# Rescue of aged muscle stem cell intrinsic quiescence defects by AKT inhibition revealed with a 3D biomimetic culture assay

**DOI:** 10.1101/2022.06.15.496252

**Authors:** Erik Jacques, Yinni Kuang, Allison P. Kann, Robert S. Krauss, Penney M. Gilbert

## Abstract

Adult skeletal muscle harbors a population of muscle stem cells (MuSCs) that are required to repair or reform multinucleated myofibers after a tissue injury. In youth, a portion of MuSCs return to a reversible state of cell cycle arrest termed ‘quiescence’ after injury resolution. By contrast, a proportion of aged MuSCs exist in a semi-activated state under homeostatic conditions, and prematurely respond to subsequent injury cues, thereby failing to return the tissue to its pre-injury state. The heterogeneity of MuSC function is linked to quiescence depth, but regulation of the balance between MuSC quiescence and activation in youth and in age is incompletely understood. This is due in part to the paucity of scalable methods that support MuSC quiescence in culture, and in turn necessitates reliance on low-throughput *in vivo* studies. To fill this gap, we developed a simple, 96-well format method to inactivate MuSCs isolated from skeletal muscle tissue, and return them to a quiescent-like state for at least one-week by culturing them within a three-dimensional engineered sheet of myotubes. Seeding the myotube sheets with different numbers of MuSCs elicited population-level adaptation activities that converged on a common steady-state niche repopulation density. By evaluating MuSC engraftment over time in culture, we observed reversible cell cycle exit that required both myotubes and a 3D culture environment. Additional quiescence-associated hallmarks were identified including a Pax7^+^CalcR^+^MyoD^-^c-FOS^-^ molecular signature, quiescent-like morphology including oval-shaped nuclei and long cytoplasmic projections with N-cadherin^+^ tips, as well as the acquisition of polarized niche markers. We further demonstrate a relationship between morphology and cell fate signature using high-content imaging and CellProfiler™-based image analysis pipelines. MuSC functional heterogeneity during engraftment was observed across all metrics tested, suggesting *in vivo*-like subpopulation activities are reflected in the assay. Notably, aged MuSCs introduced into young 3D myotube cultures displayed aberrant proliferative activities, delayed inactivation kinetics, and activation-associated morphologies that we show are rescued by wortmannin treatment. Thus, this miniaturized, biomimetic culture assay offers an unprecedented opportunity to uncover regulators of quiescence in youth and in age.

## Introduction

Muscle stem cells (MuSCs) are an adult stem cell population identifiable by the selective expression of the paired-box transcription factor Pax7 in skeletal muscle tissue, and are essential to muscle development and regeneration.^1–5^ At rest, MuSCs exist in a reversible state of quiescence characterized, among others, by the absence of cell cycle indicators^6,7^ lowered metabolic activity^8^, RNA content^9^, increased expression of genes such as CalcR, CD34, Spry 1, and Sdc4^9–11^, and an elaborate morphology^12^. Anatomically, they reside between a myofiber and the surrounding basal lamina, a highly specialized microenvironment or “niche”, that conveys unto them the popularized term ‘satellite cell’.^13^ Though quiescent, they are not dormant but are in fact idling; constantly communicating with their niche and waiting to respond to stressors.^14,15^ Examples include significant physical activity causing mechanically induced damage, trauma, or exposure to myotoxic compounds that induce myofiber degradation.^16,17^ In these situations, MuSCs rapidly shift to an activated state wherein they enter cell cycle, and proliferate to produce progeny that differentiate to repair or create new myofibers, or they undertake self-renewing divisions where a subpopulation eventually return to quiescence and repopulate the niche.^18^

Quiescence is essential to ensure the long-term stability of the MuSC pool and activation is necessary to ensure the repair process.^19^ However, how the quiescence state and the process of MuSC inactivation is regulated remains largely unexplored. MuSCs are increasingly regarded as existing individually along a quiescence-activation spectrum where shifts occur during different stages of regeneration.^19^ Indeed, the depth of quiescence shows to be positively correlated with stem cell potency, or ‘stemness’, which also remains unexplained. Consequently, instances where depth of quiescence is lost, such as in aging, leads to less efficient and incomplete regeneration, and a progressive decline in MuSC number.^20^

Tissue dissection, enzymatic digestion and cell sorting imparts an injury-associated stress response to MuSCs, and causing isolation-induced activation.^21^ Therefore, studies of quiescence *in vitro* must override activation to reinstate a quiescent state. Indeed, several studies describe *in vitro* treatments to delay activation for days (at most) through manipulation of the substrate or culture media.^22–24^ Though temporary, these strategies offer a window of opportunity to study quiescence regulation, while also offering ways to augment MuSC cell-centered therapies by improving regenerative potency by conferring a quiescent state to the population.^9,22–24^ Combining a chemically defined ‘quiescence media’ with engineered muscle fibers was reported to maintain MuSCs in culture with limited proliferative activity or changes to cell volume, and sustained CD34 expression for a 3.5-day period.^11^ More recently, three-dimensional (3D) skeletal muscle macrotissue platforms were shown to support Pax7^+^ reserve cells^25,26^ within human myoblast populations to take on a reversible quiescent-like state.^27–32^ To date, a strategy to inactivate freshly isolated MuSCs in culture for >3.5 days, while supporting molecular and morphological hallmarks of quiescence has yet to be reported.

We previously reported a method to study skeletal muscle endogenous repair “in a dish” in a 24-well format by introducing MuSCs into thin sheets of engineered muscle tissue that we then injured using myotoxins.^33^ In our uninjured control tissues, we observed a non-negligible proportion of the engrafted cells remained mononucleated at the assay endpoint, in spite of the differentiation-inducing culture media used. From this, we hypothesized that the muscle tissues were providing a pro-quiescence niche. To test this, we produced miniaturized (96-well format) muscle tissues, derived from primary mouse myoblasts, into which we introduced freshly sorted mouse MuSCs. We report that within these biomimetic niches, the MuSCs rapidly inactivated for at least 7 days. Analysis of MuSC activities reflected functional heterogeneity and population level adaptations to achieve a steady-state stem cell pool size. MuSC interactions with the 3D engineered myotube niche were sufficient for inducing *in vivo*-like hallmarks of quiescence never before reported *in vitro*, including elongated nuclei and elaborated cytoplasmic projections.^12^ Integrating the culture assay with a high content imaging system and CellProfiler™-based image analysis pipelines allowed us to relate cell fate signatures to morphometric features and produced criteria to identify quiescent MuSCs based solely on morphology. Further, aged MuSCs introduced into the assay displayed phenotypic and functional defects that were rescued by wortmannin, a treatment shown by others to push activated young MuSCs into a deep quiescent state. Thus, we present a new MuSC quiescence assay that recapitulates hallmarks of young and aged homeostatic muscle “in a dish” for the first time, which enabled the identification of a previously unreported strategy to correct aged MuSC dysfunction.

## Results

### Engineered myotube templates derived from primary mouse myoblasts maintain integrity for 2-weeks in culture

We first set out to engineer a skeletal muscle microenvironment suited to investigate the ability of a 3D myotube niche to induce a quiescent-like phenotype upon freshly isolated (i.e. activated) MuSCs cultured *in vitro*. We previously reported a method to prepare thin sheets of human myotubes situated within a 24-well format, together with a strategy to evaluate mouse MuSC endogenous repair ‘in a dish’.^33^ Herein we adapted and extended the method to create thin sheets of murine myotubes that fit within a 96-well plate footprint. Briefly, we incorporated primary mouse myoblasts within a mixture of media, fibrinogen, and Geltrex™ (i.e. reconstituted basement membrane proteins). The resultant slurry was pipetted into pieces of thin, porous cellulose teabag paper, pre-adsorbed with thrombin, and situated within a 96-well plate (**Figure 1A**). In this way, fibrin hydrogel gelation is delayed until the cell/fibrinogen slurry diffuses within the thrombin-containing cellulose scaffold. Following a two-day equilibration period in growth media (GM), the tissues were transitioned to a low-mitogen differentiation media (DM) to support multinucleate myotube formation within the cellulose reinforced fibrin hydrogel (**Figure 1A-C**).

**Figure 1.**
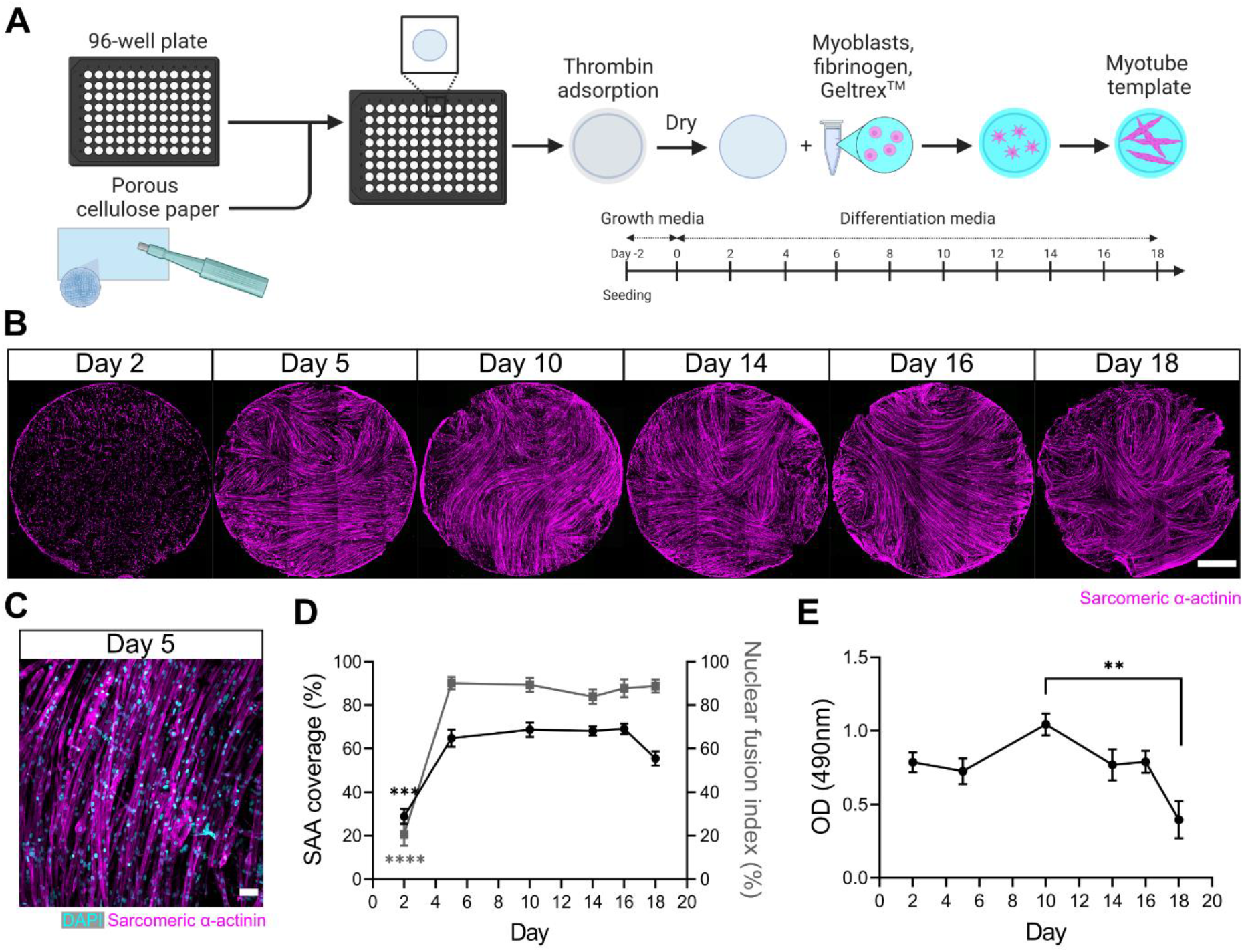
A 3D murine skeletal muscle myotube template with a 96-well footprint. **(A)** Schematic overview of the strategy used to generate myotube templates with an associated timeline for downstream culture (made with BioRender). **(B)** Representative confocal stitched images of myotube templates labelled for sarcomeric α-actinin (SAA) (magenta) at days 2, 5, 10, 14, 16, and 18 of culture. Scale bar, 1 mm. **(C)** Representative confocal image of myotubes at day 5 labelled with DAPI (cyan) and SAA (magenta). Scale bar, 50 µm. **(D)** Quantification of SAA area coverage (left-axis) and nuclear fusion index (right-axis) of myotube templates at days 2, 5, 10, 14, 16, and 18 of culture. n=9-16 across N=3-6 independent biological replicates. Graph displays mean ± s.e.m.; one-way ANOVA with Tukey post-test, minimum *** p=0.002 (SAA coverage) **** p<0.0001 (nuclear fusion index). **(E)** Optical density (OD) at 490 nm of media after myotube template incubation with MTS assay reagent on days 2, 5, 10, 14, 16, and 18 of culture. n=9-12 across N=3-4 independent biological replicates. Graph displays mean ± s.e.m.; one-way ANOVA with Tukey post-test, ** p=0.0033.

Spontaneous twitch contractions were first observed on day 4 (data not shown). Peak myotube content (≈65% by sarcomeric α-actinin (SAA) tissue coverage) and a nuclear fusion index of 90% was achieved by 5 days in DM with as few as 25,000 cells per tissue (**Figure 1B-D, Supplementary Figure 1**). Since myotube degradation could serve as an activation cue for the engrafted MuSCs, we evaluated the integrity of the tissues over time in culture. Starting on day 18, a visual inspection of tissues revealed loss of myotubes around the periphery of the tissues and quantification of SAA coverage showed a corresponding drop (**Figure 1B,D**). A colorimetric metabolic activity assay (i.e., MTS) revealed that mitochondrial activity was significantly reduced on day 18 when compared to day 10 (**Figure 1E**), another indication that the integrity of the tissues becomes compromised at these time-points. Based on these analyses, we established conditions to engineer a mouse myotube template and concluded that day 5 to < day 18 of myotube template culture would serve as the assay window.

### MuSC populations persist in myotube templates

The engineered mouse myotube template incorporates key cellular, biochemical and biophysical aspects of the MuSC niche: myofibers and extracellular matrix (ECM).^13,34,35^ Thus, we next sought to determine whether adult mouse MuSCs could persist, in terms of pool size and Pax7 expression, when introduced to these biomimetic cultures. Firstly, we adapted a magnetic-activated cell sorting (MACS) protocol as a convenient and fast alternative to fluorescence-activated cell-sorting (FACS) for enriching the Pax7^+^ mononucleated cell population from digested skeletal muscle. By conducting 2 rounds of microbead based lineage depletion followed by integrin α-7 enrichment, we achieved an average purity of 93% Pax7^+^ cells (**Supplementary Figure 2**), which meets FACS purity values reported by others.^36,37^ Using this protocol, Pax7^+^ MuSCs were enriched from the enzymatically dissociated hindlimb muscles of 129-Tg(CAG-EYFP)7AC5Nagy/J transgenic mice.^38^ Freshly sorted MuSCs were seeded onto day 5 myotube templates, and the tissue co-cultures processed for analysis at 1, 3, and 7 days post-engraftment (DPE) (**Figure 2A**). Over the one-week culture period, the Pax7^+^ mononuclear donor (YFP^+^) cells were seen distributed throughout the myotube template and adopting an elongated morphology that aligned with the local myotubes (**Figure 2B-C**). We investigated the effect of introducnig different numbers of MuSCs onto individual myotube templates, by quantifying the population of Pax7^+^ mononuclear donor cells over time. Seeding 500 MuSCs resulted in a relatively stable pool size over time. Interestingly, when a higher (1500 or 2500) or lower number of MuSCs were introduced to myotube templates, over time the number of Pax7^+^ donor cells converged to match the pool size attained in the 500 MuSC condition (**Figure 2D**). Collectively, these data indicate that the engrafted MuSC population persists and establishes a steady-state cell density within the engineered niche.

**Figure 2.**
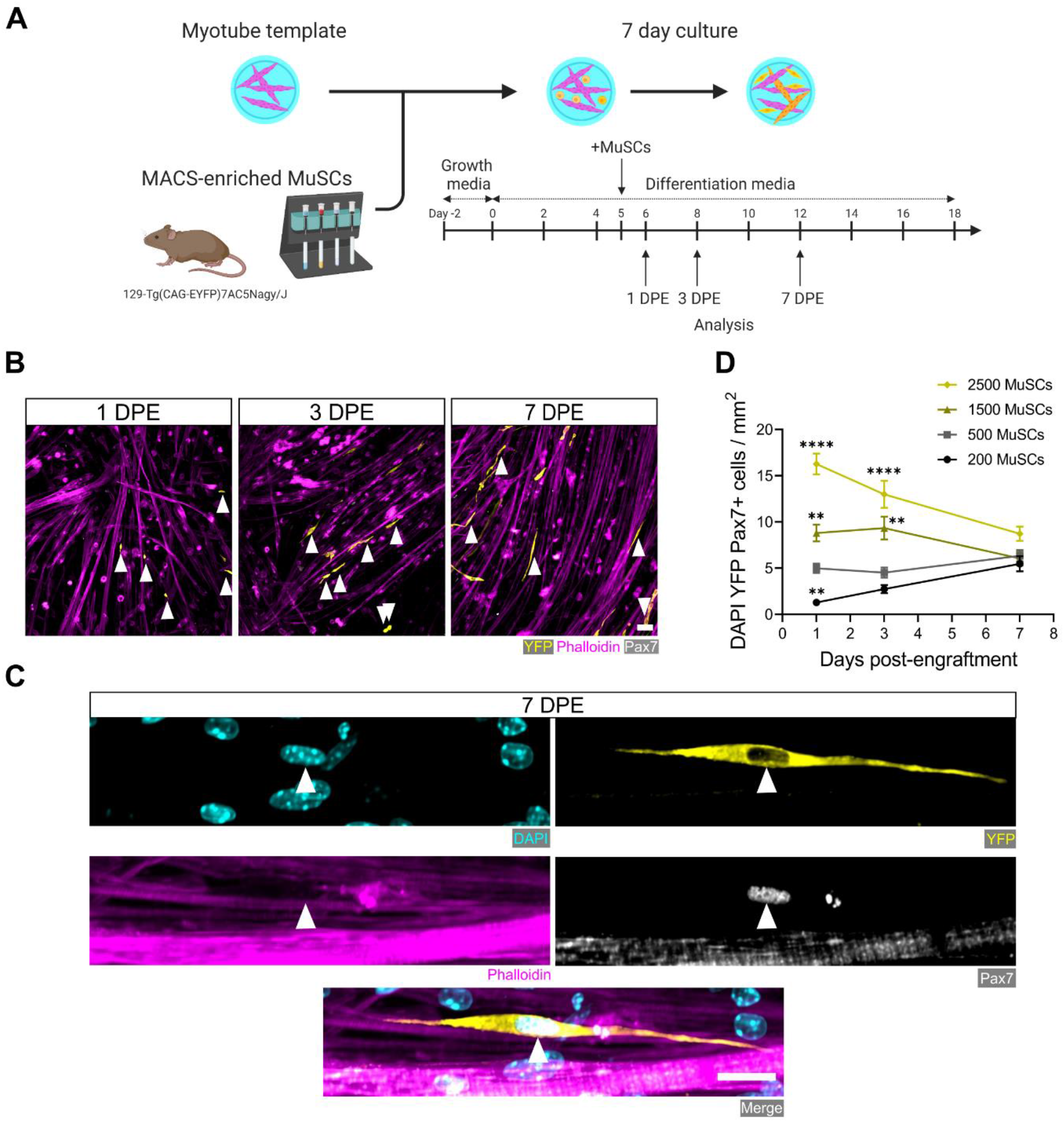
Engrafted MuSCs persist in myotube template cultures and achieve a steady-state population density. **(A)** Schematic overview of the engraftment of freshly isolated MuSCs and the timeline for downstream analysis (made with BioRender). **(B)** Representative confocal images of myotube templates (phalloidin: magenta) with engrafted MuSCs (YFP: yellow, Pax7: white, white arrows) at 1, 3 and 7 days post-engraftment (DPE). Scale bar, 50 µm. **(C)** Representative confocal image of a donor MuSC (DAPI: cyan, YFP: yellow, Pax7: white) indicated with a white arrow, and myotubes (phalloidin: magenta) at 7 DPE. Scale bar, 20 µm. **(D)** Quantification of mononuclear DAPI^+^YFP^+^Pax7^+^ cell density per mm^2^ at 1, 3 and 7 DPE across different starting MuSC engraftment numbers (200, 500, 1500, and 2500). n=9-15 across N=3-5 independent biological replicates. Graph displays mean ± s.e.m.; one-way ANOVA with Dunnet test for each individual timepoint comparing against the 500 MuSC condition, ** p=0.0025, 0.0051, 0.0029 **** p<0.0001.

### MuSCs reversibly inactivate within myotubes templates

We next studied the behavior and fate of freshly isolated MuSCs engrafted within myotube templates and determined that they inactivate over a 7-day culture period, and can be coaxed to reactivate by injury stimuli. We began by evaluating MuSCs within the engraftment condition that lent to a stable population density over time (i.e. 500 MuSCs per tissue). Calcitonin receptor (CalcR) expression is a hallmark of quiescent MuSCs.^39–42^ Indeed, at the protein level, CalcR is expressed by quiescent MuSCs, but is then absent from all MuSCs within 48-hours of an *in vivo* myotoxin injury or within 48-hours of prospective isolation followed by *in vitro* culture.^39,40,42^ In the context of our 3D culture assay, the majority of MuSCs expressed CalcR at 1 DPE, with a sharp decline in the proportion of CalcR^+^ donor cells observed at 3 DPE (**Supplementary Figure 3)**. Interestingly, ∼15% of donor MuSCs were CalcR^+^ at both 3 and 7 DPE (**Supplementary Figure 3**). Given the lack of evidence for CalcR^+^ MuSCs in prolonged in vitro cultures, we posited that this sub-population might be reflective of MuSCs that had resisted activation in favour of maintaining a more quiescent-like state, which we sought to interrogate further.

After a single day of culture, we found that ≈75% of the donor MuSCs (YFP^+^Caveolin-1^+^ cells) engrafted within the myotube templates expressed the transcription factor c-FOS, among the earliest transcriptional events reported to-date in the MuSC activation sequence.^9,43–45^ The existence of c-FOS^-^ donor cells at this time-point is consistent with the notion of an activation refractory sub-population. By 3 DPE, the proportion of caveolin-1^+^c-FOS^+^ mononuclear cells dropped to ≈30%, with similar proportions observed on 7 DPE (**Figure 3A-B**). The maintenance of a steady-state population of donor MuSCs from 1 DPE to 3 DPE, coupled with the rapid loss of c-FOS immunolabeling by 3 DPE, suggests that myotube template culture induces MuSCs to inactivate.

**Figure 3.**
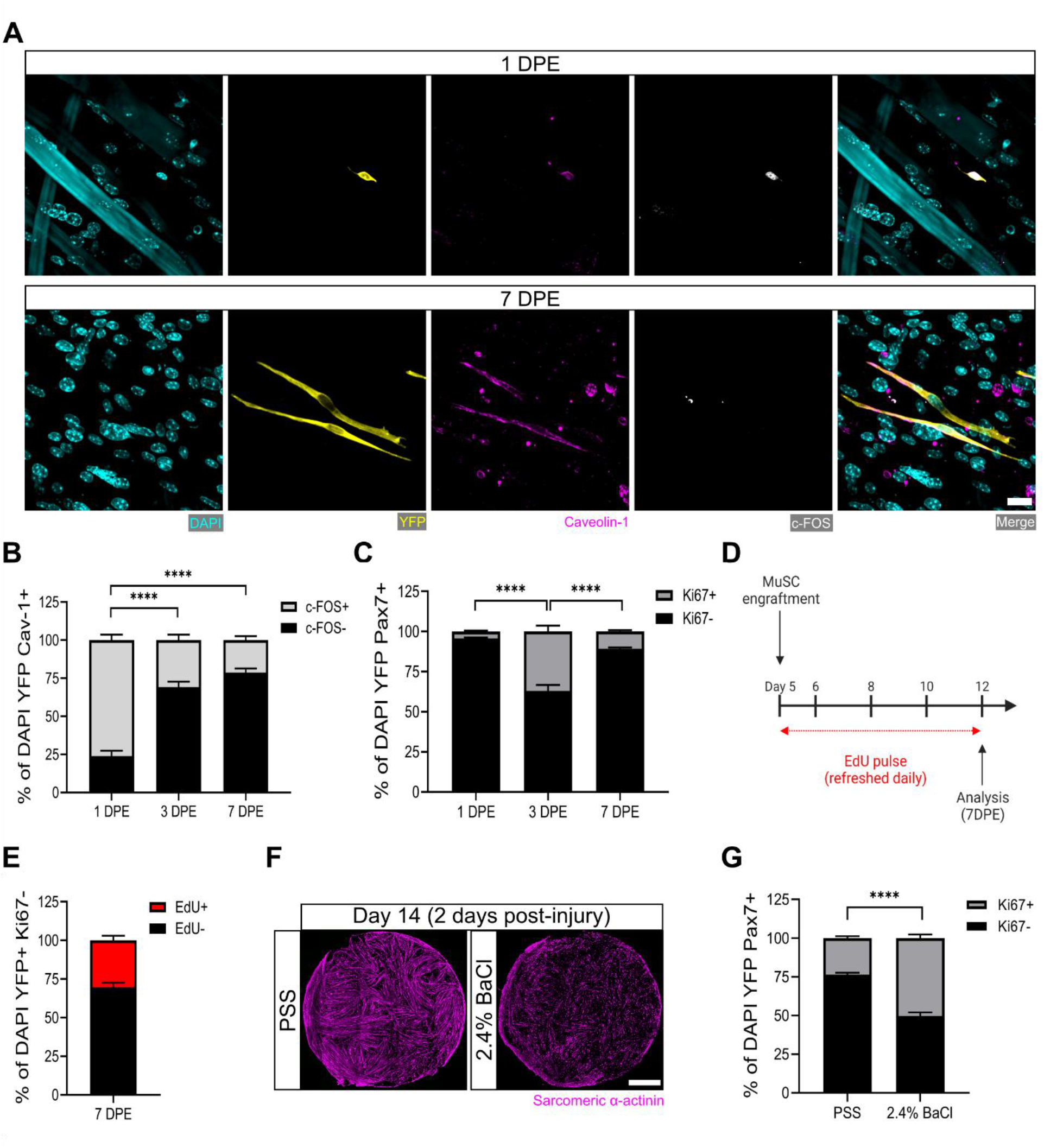
MuSCs engrafted within engineered muscle tissue exit cell-cycle and inactivate. **(A)** Representative confocal image of a mononuclear cell (DAPI: cyan) positive for YFP (yellow), caveolin-1 (magenta) and c-FOS (white) at 1 DPE (Top), and a c-FOS^-^ cell at 7 DPE (Bottom). Scale bar, 20 µm. **(B)** Stacked bar graph showing proportions of c-FOS+/**-** cells at 1, 3 and 7 DPE in the DAPI^+^YFP^+^Cav-1^+^ population. n=9 across N=3 independent biological replicates. Graph displays mean ± s.e.m. for c-FOS^+^ and c-FOS^-^; one-way ANOVA with Tukey post-test comparing the FOS^-^ proportions of each timepoint, **** p<0.0001. **(C)** Stacked bar graph showing proportions of Ki67+/- cells at 1, 3 and 7 DPE in the DAPI^+^YFP^+^Pax7^+^ population. n=10-11 across N=3-4 independent biological replicates. Graph displays mean ± s.e.m. for Ki67^+^ and Ki67^-^; one-way ANOVA with Tukey post-test comparing the Ki67^-^ proportions of each timepoint, **** p<0.000.1 **(D)** Timeline of EdU/Ki67 co-labelling experiment (made with BioRender). **(E)** Stacked bar graph showing proportions of EdU+/- cells at 7 DPE in the DAPI^+^YFP^+^Ki67^-^ mononuclear cell population. n=15 across N=5 independent biological replicates. Graph displays mean ± s.e.m. for EdU^+^ and EdU^-^. **(F)** Representative confocal stitched images of myotube templates (SAA: magenta) 2 days after a 4-hour exposure to the physiological salt solution (PSS) control or a 2.4 % barium chloride (BaCl_2_) solution. Scale bar, 1 mm. **(G)** Proportion of Ki67+/- cells at 2 DPI in the DAP^+^YFP^+^Pax7^+^ population. n=16, 18 across N=5, 6 biological replicates. Graph displays mean ± s.e.m. for Ki67^+^ and Ki67^-^; unpaired t-test of the Ki67^-^ proportions of both conditions, **** p<0.0001

Consistently, when we quantified the incidence of MuSCs in the active phase of the cell cycle via Ki67 labelling, we found that at 3 DPE, only ≈1/3 of the Pax7^+^ mononuclear donor cell population was Ki67^+^, and this dropped to ≈10% by 7 DPE (**Figure 3C**). To better resolve the proliferative trajectory of the engrafting MuSCs, we conducted a Ki67 co-labelling study whereby 5-ethanyl-2’-deoxyuridine (EdU) was refreshed in the culture media daily over the 1-week culture period (**Figure 3D**). Of the Ki67^-^ mononuclear donor cells present at 7 DPE, the vast majority were EdU^-^ (**Figure 3E**). ≈30% were EdU^+^, indicating cell cycle entry at some point during the one-week culture period, and a cessation by 7 DPE (**Figure 3E**). This correlates well with the proportion of Ki67^+^ MuSCs we observed at 3 DPE (**Figure 3B**). This data, together with a scarcity of EdU^+^ myonuclei observed in the cultures (data not shown), suggests that the main fate of the EdU labeled MuSCs is eventual cell-cycle exit, and not myotube fusion.

Lastly, we sought to understand whether the inactivated donor cells at 7 DPE were capable of re-entering the cell-cycle. We first established a barium chloride exposure protocol that induced effective clearing of the myotubes with a non-significant change to MuSC population density (**Figure 3F** and **Supplementary Figure 4**). We then analyzed the mononuclear YFP^+^Pax7^+^ population 2 days post-injury and observed a statistically significant increase in the proportion of Ki67^+^ cells as compared to the control condition (**Figure 3G**). Thus, myotube template cultures allow for inactivation and cell-cycle exit of engrafted MuSCs, which can be reversed with the injury-associated stimuli provided by barium chloride exposure.

### Engrafted MuSCs adapt their pool size to a myotube template threshold

Regardless of the initial size of the MuSC pool, a common mononuclear YFP^+^Pax7^+^ cell density was attained by 7 DPE (**Figure 2D**). To uncover cellular mechanisms underlying the acquisition of a MuSC steady-state population density, we investigated how the donor MuSC pool responded under a set of distinct starting conditions. We began by extending the EdU/Ki67 co-labelling study (**Figure 3D-E**) to include an evaluation of conditions where more (1500, 2500) or less (200) MuSCs were seeded onto the myotube templates. Compared with the 500 MuSC seeding condition, we found a significant increase in the proportion of mononuclear YFP^+^Ki67^-^ cells that were EdU^+^ at 7 DPE in cultures seeded with 200 MuSCs, suggesting the MuSC pool expanded to attain a steady-state density (**Supplementary Figure 5A,C**). By contrast, in conditions where >500 MuSCs were seeded, a significant decrease in the proportion of mononuclear YFP^+^Ki67^-^ cells that were EdU^+^ at 7 DPE was observed (**Supplementary Figure 5C**). In these conditions, a decrease in the MuSC pool size by 7 DPE could be achieved through cell death or by fusion into myotubes. Consistent with the latter hypothesis, upon visual inspection we saw a qualitatively greater number of donor derived myotubes in the cultures seeded with >500 MuSCs (**Supplementary Figure 5B**), which was confirmed by quantifying the percentage area of myotube templates covered by YFP signal (**Supplementary Figure 5D**). In sum, we conclude that MuSCs meet a steady-state population density via increased proliferation when beginning below the 500-cell threshold, and with increased cell fusion when beginning above it.

### A three-dimensional myotube culture is required for a persistent MuSC population *in vitro*

The rapid inactivation and subsequent maintenance of Pax7^+^ MuSCs engrafted within the 3D myotube templates (**Figures 2-3**) represents a divergent phenotype when compared to conventional 2D culture (**Figure 4A-B**).^46^ Therefore, we next sought to elucidate culture design criteria that served to support MuSC inactivation and pool maintenance over time. We first explored the response of MuSCs seeded onto tissues on Day 0 of myotube template differentiation, a time-point corresponding to the earliest myocyte fusion events, and therefore when myotubes were absent from the tissues. Compared to MuSCs seeded on myotube templates on Day 5 of differentiation, Day 0 seeding resulted in a progressive loss of YFP^+^Pax7^+^ mononuclear cells, and most of those that remained were Ki67^+^ (**Figure 4C-D**). The striking contrast in YFP^+^ myotube content observed at 7 DPE upon comparing these two conditions suggests that the MuSCs engrafted on Day 0 had undergone differentiation (**Supplementary Figure 6**). We next determined whether a 3D cellulose-reinforced hydrogel alone was sufficient to support MuSC inactivation and maintenance, since the myocytes present on Day 0 of differentiation may have exerted a dominant effect overriding contributions of the 3D culture environment. However, this notion was abandoned upon finding that the outcome of this culture scenario (**Figure 4E**) very closely matched what we observed when the MuSCs were cultured in 2D Geltrex-coated culture wells (**Figure 4B**); a loss of the YFP^+^Pax7^+^ mononuclear population over time. Our results instead seemed to suggest that the myotube template played a central role in inactivating and maintaining a persistent population of MuSCs in culture. Indeed, upon adding MuSCs to a Day 5 monolayer of myotubes in 2D culture, a Pax7^+^ population was maintained over the one-week culture period (**Figure 4F**). However, in striking contrast to the 3D myotube template culture (**Figure 4C**), only a minority of the Pax7^+^ donor cells were Ki67^-^ at 7 DPE (**Figure 4F**). From this iterative analysis, we conclude that myotubes are necessary for Pax7^+^ MuSC persistence, and that the combination of myotubes and a three-dimensional culture environment drives the MuSC inactivation process.

**Figure 4.**
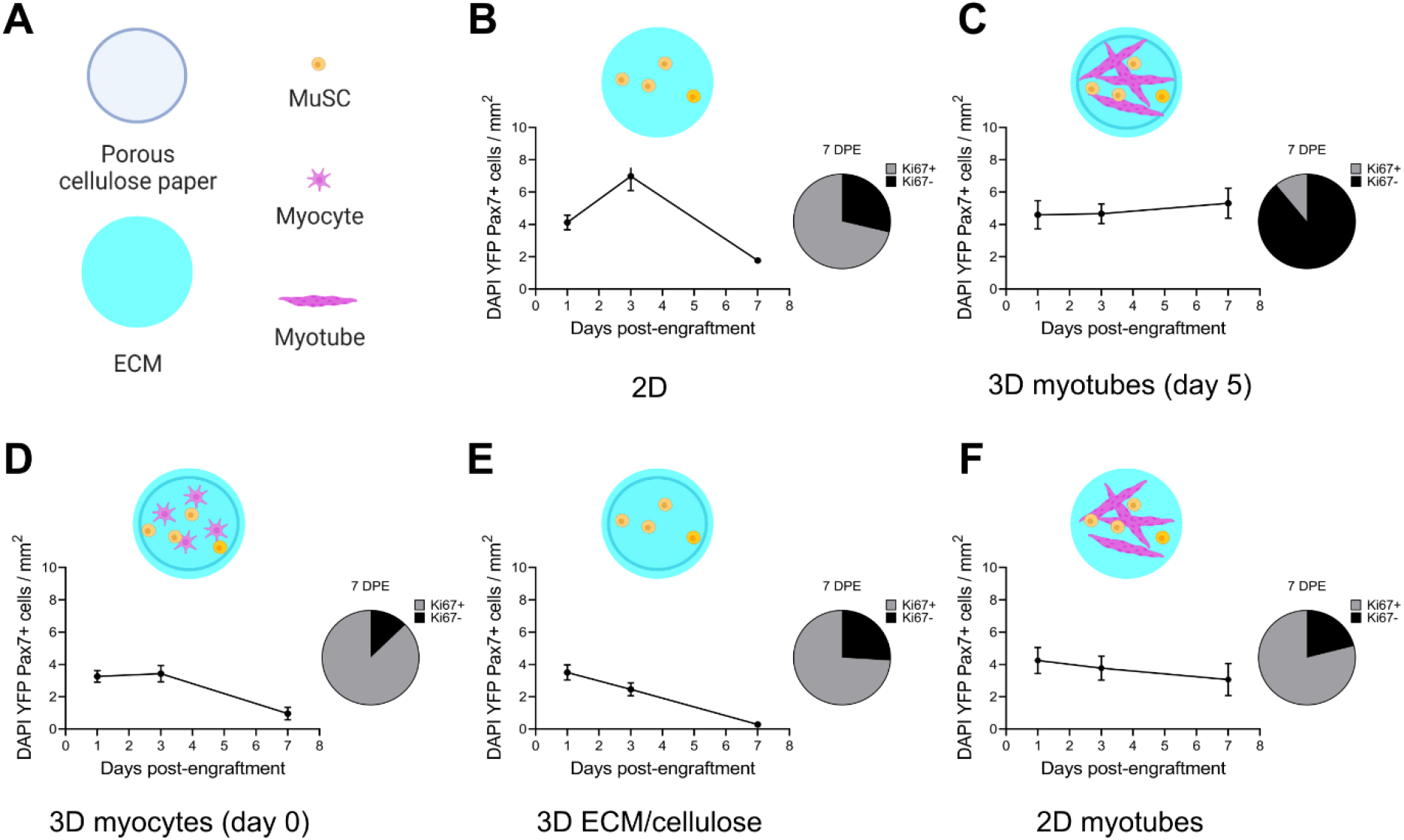
Permissive culture conditions for a persistent MuSC population *in vitro*. **(A)** Key for figure icons. **(B-F)** Line graphs of mononucleated DAPI^+^YFP^+^Pax7^+^ cell density at 1, 3 and 7 DPE (left) and pie charts showing the proportion of Ki67+/- cells at 7 DPE (right) for cells seeded into a 2D microwell with a Geltrex™ coating **(B)**, engrafted into 3D myotube templates on day 5 **(C)** vs day 0 **(D)** of differentiation. Additional comparisons include engraftment into a 3D cellulose reinforced extracellular matrix (ECM) hydrogel on day 5 **(E)**, or onto a 2D monolayer of myotubes with a Geltrex™ undercoating on day 5 of differentiation **(F)**. n=6-9 from N=2-3 independent biological replicates. Graphs display mean ± s.e.m.

### Engrafted MuSCs adopt quiescent-like morphologies that predict cell fate signature

Qualitatively, the engrafted MuSCs in our cultures adopted an elongated morphology over time (**Figure 2B-C, Figure 3A, Figure 5A**), reminiscent of quiescent MuSCs *in vivo*, and contrasting against with morphologies observed in 2D cultures (**Supplementary Figure 7**).^12,47,48^ Therefore, we next overcame a significant data analysis bottleneck by establishing and validating a CellProfiler™-based image analysis pipeline in order to segment and evaluate donor MuSCs in our phenotypic datasets (see Methods and **Supplementary Figure 8**).^49^ The cytoplasmic elongation of mononucleated Pax7^+^ donor cell bodies was captured by applying a ratio of max/min feret diameter to segmented images of tissues immunostained for YFP, Pax7, and DAPI. The roundness of nuclei within mononucleated YFP^+^Pax7^+^ cells was evaluated using the measurement of eccentricity, whereby a value of 0 corresponds to a perfect circle, and a value of 1 to a straight line (**Figure 5B**). With this pipeline, we determined that the Pax7^+^ donor cell population progressively shifted from low max/min feret diameter ratios and eccentricities (lower left quadrant) to high max/min feret diameter ratios and eccentricities (upper right quadrant) over time in 3D myotube culture (**Figure 5C**). The rice-like nuclear morphology and elaborated cytoplasmic projections of the Pax7^+^ donor cells at 7 DPE resembled quiescent features of MuSCs *in vivo* that were shown to be induced and maintained by tipping the Rho family GTPase balance to favour cytoskeletal remodelling events caused by Rac signaling.^48^

**Figure 5.**
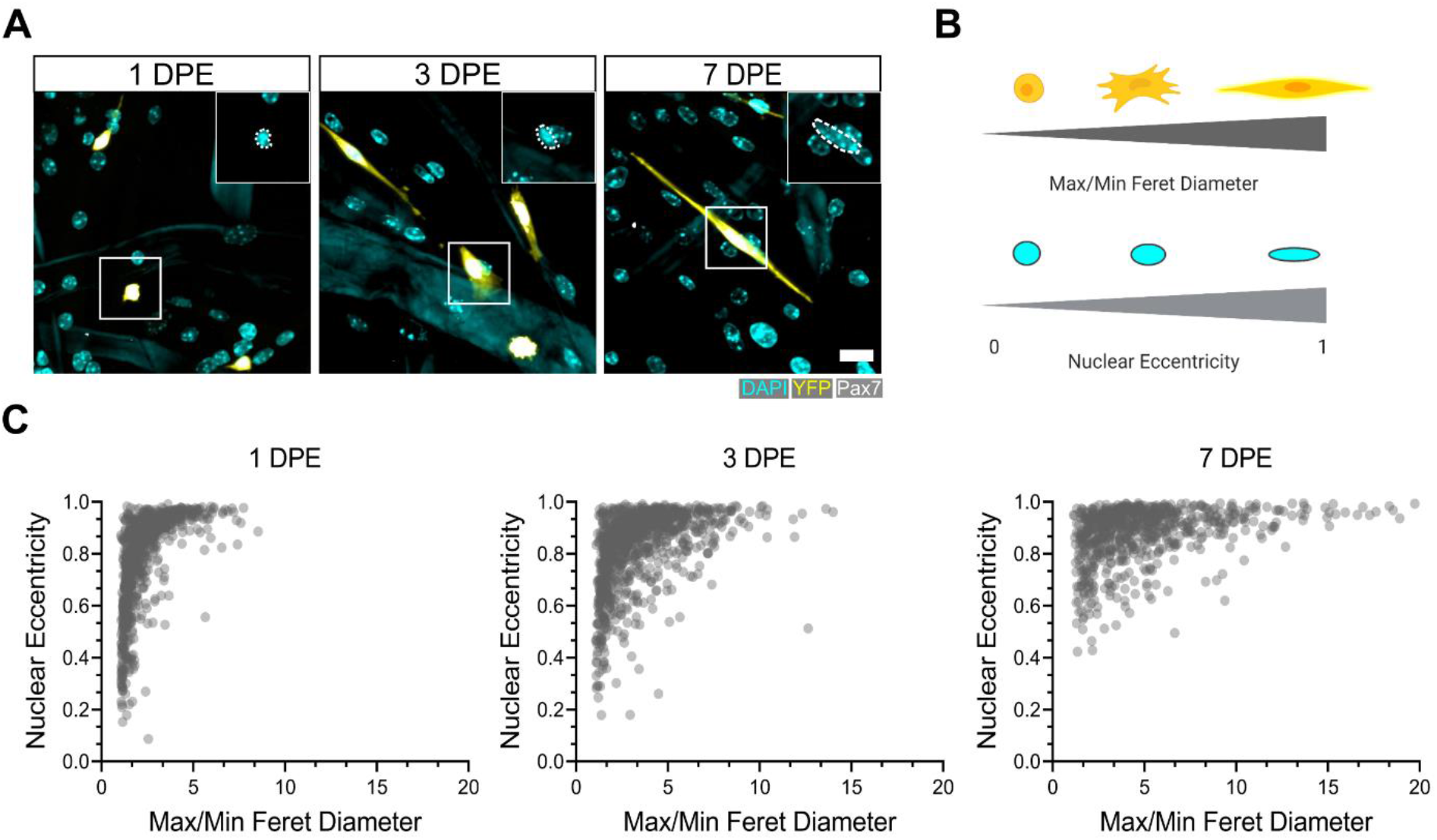
Morphological evolution of engrafted MuSCs. **(A)** Representative confocal images of MuSCs (DAPI: cyan, YFP: yellow, Pax7: white) with distinct morphological features at 1, 3 and 7 DPE. Insets highlight nuclear morphology with a white dotted outline. Scale bar, 20 µm. **(B)** Schematic demonstrating the morphological features quantified using CellProfiler™ (made with BioRender). **(C)** Dot plot graphs showing individual Pax7^+^ donor cells and their associated max/min feret diameter ratio and nuclear eccentricity at 1 (left), 3 (middle), and 7 DPE (right). n=916, 980 and 737 across N=3-4 biological replicates.

We next sought to determine whether the donor MuSC morphologies observed in our cultures corroborated with the activation status of the cells. We introduced immunolabelling for MyoD, which, together with Pax7 staining, delivered molecular signatures for activated (Pax7^+^MyoD^+^) and inactivated (Pax7^+^MyoD^-^) donor cell populations. As expected, the ratio of Pax7^+^MyoD^+^ to Pax7^+^MyoD^-^ donor cells over time followed a trend similar to Ki67 status (**Figure 3C**), with a transient increase in Pax7^+^MyoD^+^ cells at 3 DPE and a predominance of Pax7^+^MyoD^-^ cells at 7 DPE (**Supplementary Figure 9A-B**). By evaluating the mean max/min feret diameter ratio and mean eccentricity values of the Pax7^+^MyoD^+^ and Pax7^+^MyoD^-^ cell populations, we found that nuclear eccentricity differs between the populations by 3 DPE, while population divergence according to max/min feret diameter ratio (>2-fold) emerged a bit later, at 7 DPE (**Supplementary Figure 9C**). Specifically, nuclear morphology of the Pax7^+^MyoD^-^ population showed a progressive, statistically significant transition to a rice-like shape, while Pax7^+^MyoD^+^ nuclei remained more rounded. Elongation of the cell body and elaborate projections were features that exclusively characterized the Pax7^+^MyoD^-^ cell population, and emerged between 3 DPE and 7 DPE. Indeed, donor cells with a max/min feret diameter ≥5.8 uniformly displayed the Pax7^+^MyoD^-^ signature of inactivated cells (**Supplementary Figure 9A**). Taken together, morphological analysis of the engrafted MuSCs suggests that changes in nuclear morphology precede cell body extension and establishment of quiescent-like projections during the inactivation process, and that morphometric features alone may predict MuSC inactivation status.

### Engrafted MuSCs establish a polarized niche

We next evaluated additional hallmarks of quiescent MuSCs including the spatial organization of cadherins, integrins, and extracellular matrix proteins relative to their niche.^50–52^ *In vivo*, MuSCs are identified anatomically by their positioning sandwiched between a myofiber and the surrounding basal lamina.^13^ This polarized niche lends to the intracellular segregation or deposition of proteins within MuSCs to the apical side facing the myofiber (e.g. N-cadherin) or to the basal side facing the basal lamina (e.g. integrin α-7, laminin).^51,52^ By evaluating immunolabelled tissues at 7 DPE we found that most mononucleated donor cells had an elongated morphology and were closely associated with multinucleated myotubes. In more than two-thirds of donor cells, M-cadherin expression restricted to the apical interface was observed (**Supplemental Figure 10**).

It was recently discovered that quiescent MuSCs localize the N-cadherin adhesion molecule to the tips of elaborated cytoplasmic projections (coined ‘quiescent projections’) ^48^, a feature we also observed within our culture assay at 7 DPE (**Figure 6A**). While examples such as these were relatively uncommon occurrences, in each case the Pax7^+^ donor cell morphology was characterized by a long oval-shaped nucleus and very long, elaborated cytoplasmic projections. Furthermore, we identified examples of polarized distribution of M-cadherin and integrin α-7 or laminin α-2 in Pax7^+^ donor cells (**Figure 6B** and **Supplementary Figure 11**). This evidence demonstrates that the engrafted MuSCs can recapitulate anatomical hallmarks of MuSCs residing within adult homeostatic skeletal muscle, and suggests that acquisition of these features is dependent on interactions with MuSCs and their immediate myofiber niche.

**Figure 6.**
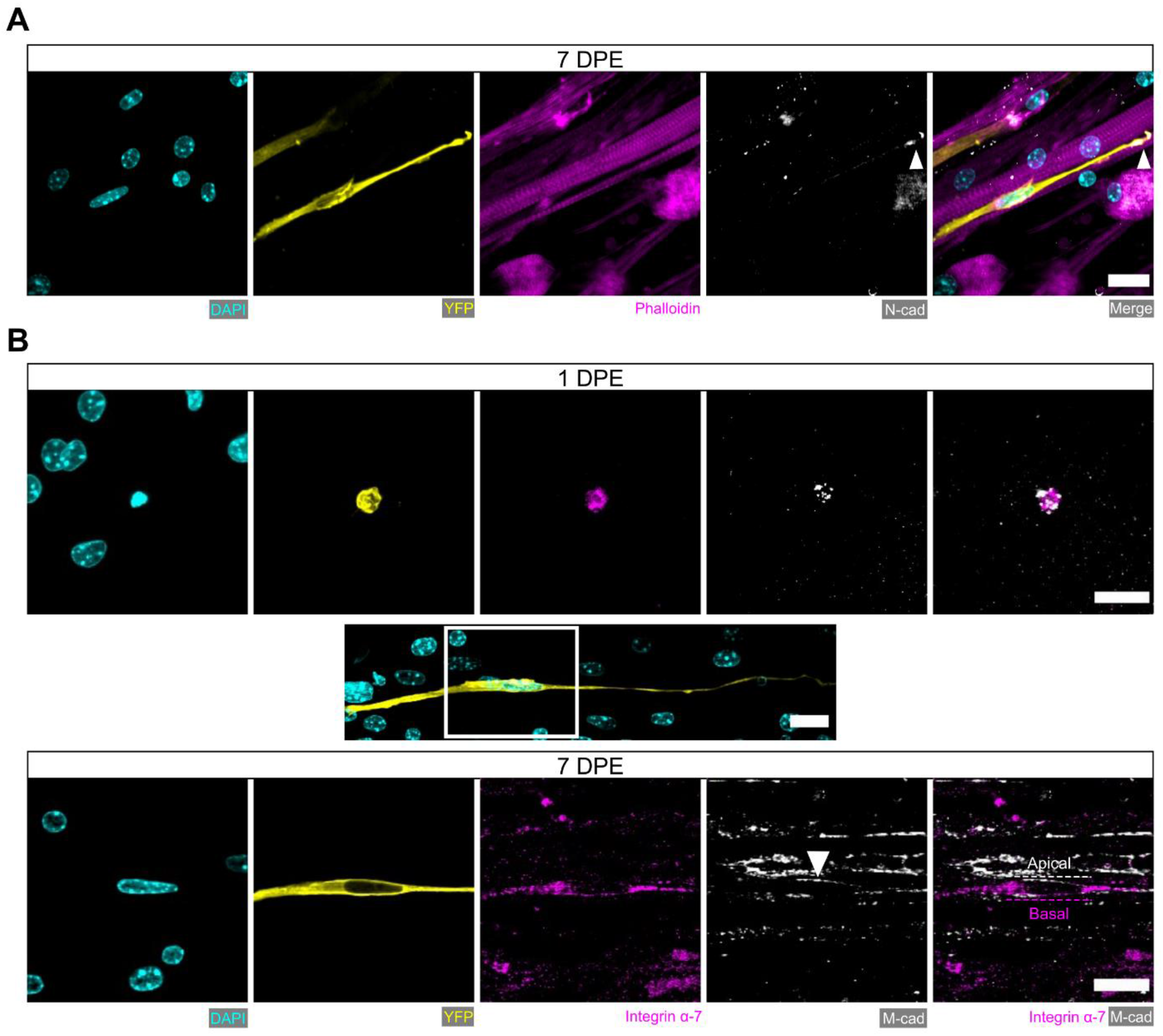
Engrafted MuSCs display quiescence and niche-related hallmarks. **(A)** Representative confocal image of a mononuclear donor cell (DAPI: cyan, YFP: yellow) with neighbouring myotubes (Phalloidin: magenta) and N-cadherin (white) localized to the tip of the donor cell projection (white arrowhead). Scale bar, 20 µm. **(B)** Representative confocal images of a mononuclear donor cell (DAPI: cyan, YFP: yellow) at 1 DPE (top) and 7 DPI (middle and bottom) expressing integrin α-7 (magenta) and M-cadherin (white). Middle inset image channels are separated to produce the bottom images to highlight the polarization of integrin α-7 and M-cadherin (white arrow) to basal and apical orientations, respectively (dotted lines). Scale bars, 20 µm.

### Aged MuSCs exhibit quiescence-related defects that can be rescued by Akt inhibition

We have shown that freshly isolated MuSCs are coaxed into a quiescent-like state that is characterized by cell cycle exit, a Pax7^+^MyoD^-^c-Fos^-^ signature, morphological and niche associated features, when introduced to a 3D myotube culture environment. A hallmark of MuSCs residing within aged muscle is precocious activation, owing to an improper maintenance and/or return to a quiescent state.^20,53–56^ Indeed, a proportion of aged MuSCs remain ‘primed’ to activate. As a result of the improper repair kinetics caused by the ‘primed’ state, these MuSCs fail to meet regenerative demand and are eventually depleted with further age.^20,57^ We next leveraged our assay to evaluate possible intrinsic defects in aged MuSC quiescence that might be apparent when they are decoupled from an aged niche environment. Upon seeding 500 MuSCs isolated from aged muscle onto a young 3D myotube template, we quantified a ≈2-fold increase in Pax7^+^ mononucleated donor cell density by 3 DPE relative to tissues engrafted by young MuSCs and analyzed at the same time-point (**Figure 7A** and **Supplementary Figure 12A**). Consistently, a greater proportion of aged as compared to young MuSCs were Ki67^+^ at 3 DPE, and aged MuSC morphology at this time-point diverged significantly from that observed of young MuSCs in 3D myotube cultures (**Figure 7B-C** and **Figure 2B**). By 7 DPE, aged MuSC engrafted cultures showed a small, but significant, decrease in population density compared to young MuSC engrafted tissues (**Figure 7A**). This, coupled with the trending increase in donor cell GFP^+^ signal covering tissues at this timepoint (**Supplementary Figure 13**), suggested that the aged MuSCs were unable to maintain pool size and that the production of Pax7^+^ donor cells observed at 3 DPE culminated in differentiation to myotubes. Nonetheless, the mononucleated aged Pax7^+^ donor cell population that persisted throughout the culture period showed a decline in the proportion of c-FOS^+^ (**Figure 7D**) and Ki67^+^ (**Figure 7C**) cells with time, albeit with delayed inactivation kinetics when compared to young MuSCs (**Figure 7C-D**). To further evaluate the quiescent-like state of the engrafted MuSC populations, we quantified the morphology of individual Pax7^+^ donor cells at 7 DPE. On average, aged donor cells had reduced max/min feret diameter ratio and nuclear eccentricity when compared to young cells at this timepoint, which correlates to the more contracted/rounded morphological characteristics of activated MuSCs (**Figure 7E-G**).

**Figure 7.**
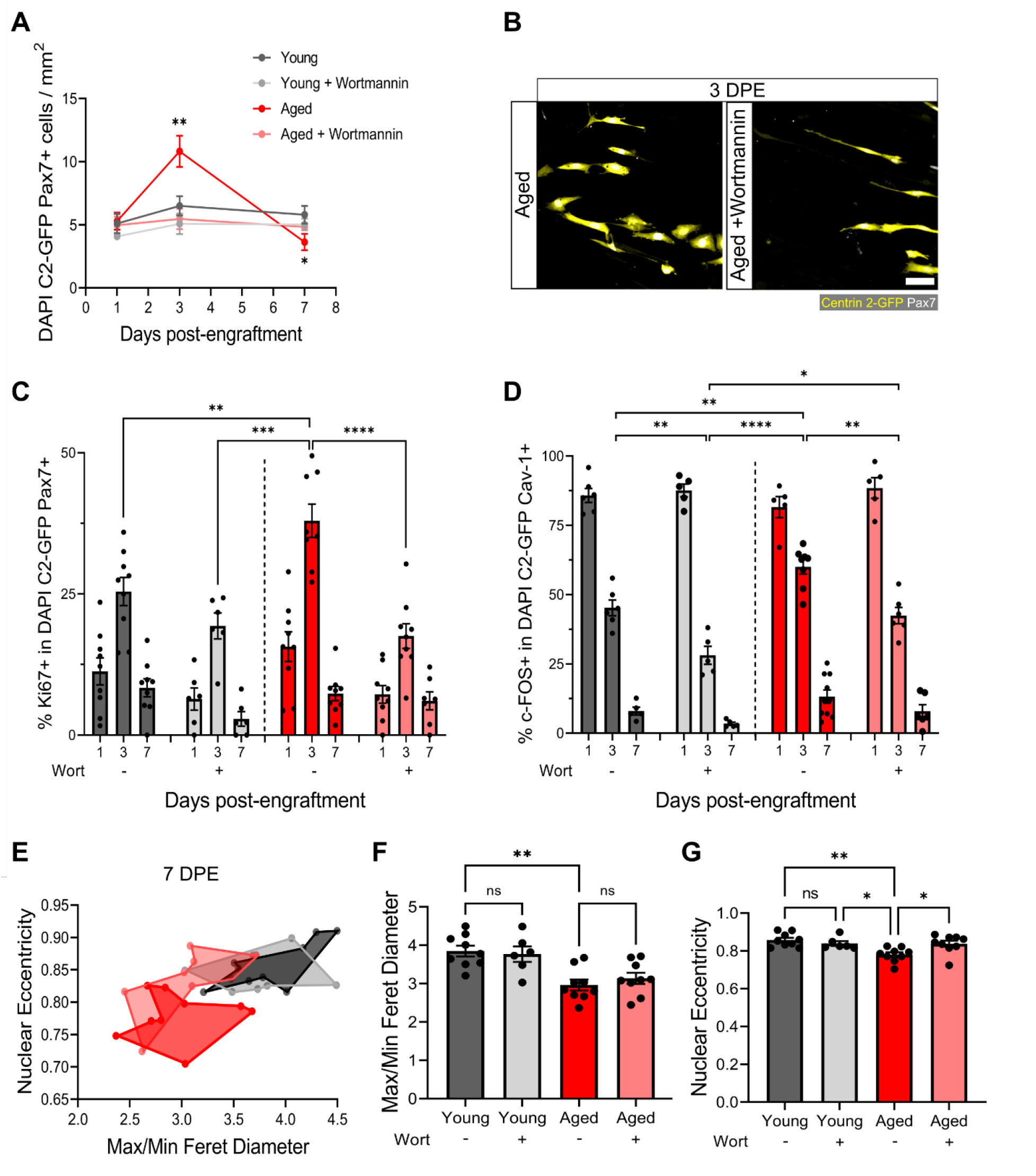
Aberrant pool size maintenance and inactivation in aged MuSCs is rescued by wortmannin. **(A)** Quantification of mononuclear DAPI^+^Centrin 2-GFP (C2-GFP)^+^Pax7^+^ cell density per mm^2^ at 1, 3 and 7 DPE between engrafted young and aged MuSCs +/- wortmannin (wort) treatment. n=6-9 across N=2-3 independent biological replicates, graph displays mean ± s.e.m.; one-way ANOVA with Dunnet test for each individual timepoint comparing against the young condition, * p=0.0262 ** p=0.0065. **(B)** Representative confocal image of donor cells (Centrin 2-GFP:yellow, Pax7:white) from the aged and aged + wortmannin conditions at 3 DPE. Scale bar, 50 µm. **(C)** Bar graph showing the percentage of Ki67^+^ cells in the DAPI^+^C2-GFP^+^Pax7^+^ mononucleated population at 1, 3 and 7 DPE across experimental conditions (young: dark grey; young + wortmannin: light grey; aged: red; aged + wortmannin: light red). n=6-9 across N=2-3 independent biological replicates, graph displays mean ± s.e.m. with individual technical replicates; one-way ANOVA with Tukey’s post-test comparing the conditions against each other at the 3 DPE timepoint, ** p= 0.0064 *** p=0.0003 **** p<0.0001 (comparisons not shown are ns). **(D)** Bar graph showing the percentage of c-FOS^+^ cells in the DAPI^+^C2-GFP^+^Cav-1^+^ mononucleated population at 1, 3 and 7 DPE across experimental conditions (young: dark grey; young + wortmannin: light grey; aged: red; aged + wortmannin: light red). n=5-10 across N=2-3 independent biological replicates, graph displays mean ± s.e.m. with individual technical replicates; one-way ANOVA with Tukey’s post-test comparing the conditions against each other at the 3 DPE timepoint, * p=0.0169 ** p= 0.0040, 0.0053, 0.0010 **** p<0.0001 (comparisons not shown are ns). **(E)** Dot graph where each dot represents the average max/min feret diameter ratio and nuclear eccentricity of the Pax7^+^ donor cells within the technical replicate (tissue) at the 7 DPE timepoint, color coded according to experimental condition (young: dark grey; young + wortmannin: light grey; aged: red; aged + wortmannin: light red). **(F)** Bar graph showing the average max/min feret diameter ratio across experimental conditions, graph displays mean ± s.e.m. with the individual technical replicates from panel E; one-way ANOVA with Tukey’s post-test, ** p=0.0010 (young vs aged + wortmannin and young + wortmannin vs aged are also ** p=0.0093, 0.0084, but not shown. All other comparisons are not significant. **(G)** Bar graph showing the average nuclear eccentricity across experimental conditions, graph displays mean ± s.e.m. with the individual technical replicates from panel (E); one-way ANOVA with Tukey’s post-test, * p=0.0402, 0.0216 ** p=0.0015. All other comparisons are not significant.

We next pursued a potential rescue of the aged MuSC phenotypes we observed in our engineered cultures. Recent work showed that FoxO transcriptions factors (TFs) are responsible for conferring a ‘genuine’ quiescent state to MuSCs, whereby genetic ablation resulted in a shift towards a ‘primed’ state.^54^ Furthermore, FoxO activity was computationally predicted to be regulated by the Igf-Akt pathway, where phosphorylated-Akt causes the phosphorylation of FoxO transcription factors and their translocation to the cytoplasm. Pharmacological inhibition of the AKT pathway using the phosphatidylinositol 3 kinase (PI3K) inhibitor, wortmannin, resulted in increased stemness in primed young MuSCs. Whether this treatment strategy is capable of rescuing aged MuSCs is currently unknown; providing an opportunity to leverage our culture model to uncover new biology. First, we confirmed that compared to young MuSCs, aged MuSCs presented increased proliferation and reduced FoxO3a nuclear fluorescent intensity in 2D culture. In this context, wortmannin treatment (10 µM) blunted cell proliferation and increased FoxO3a nuclear localization in both young and aged MuSCs (**Supplementary Figure 12A-C**). Indeed, FoxO3a nuclear fluorescent intensity was comparable between young and aged MuSCs following wortmannin treatment (**Supplementary Figure 12B**).

We then introduced wortmannin to 3D myotube cultures engrafted with young or aged MuSCs. With this treatment, the aged MuSCs maintained a stable Pax7^+^ donor population size over time, now indistinguishable from the untreated young MuSCs cultures (**Figure 7A-B**). Indeed, the proportions of Ki67^+^ and c-FOS^+^ in the aged Pax7^+^ population showed comparable kinetics to the young untreated donor MuSCs (**Figure 7C-D**). As well, the treatment encouraged a greater proportion of young MuSCs to inactivate, and with more rapid kinetics (**Figure 7C**). We also found a trending decrease in donor cell GFP^+^ signal covering tissues at 7 DPE in wortmannin treated conditions, suggesting reduced differentiation (**Supplementary Figure 13**). Finally, morphological characterization of wortmannin treated aged MuSCs at 7 DPE showed no change in average max/min feret diameter ratio, whereas we found a rescue of nuclear eccentricity that matched young MuSCs (**Figure 7E-G**). There were no shifts in the morphological profile of young MuSCs treated with wortmannin (**Figure 7E-G**).

Thus, by introducing aged MuSCs into a young myotube niche we revealed abnormal population maintenance, delays in the inactivation kinetics, and morphological features characteristic of activated MuSCs, which we show can be rescued by modulating AKT signalling, a pathway shown by others to regulate quiescence in young MuSCs.

## Discussion

We have developed an *in vitro* functional assay that rapidly induces and sustains MuSC inactivation, enabling systematic analyses of cellular and molecular mechanisms presiding over the return to quiescence for the first time. MuSCs acquired in *vivo*-like hallmarks of quiescence, that, to our knowledge have never before been reported *in vitro*. Through temporal single cell analyses, we uncovered evidence of population-level adaptations to the muscle tissue niche and functionally heterogenous MuSC sub-populations mirroring *in vivo* heterogeneous activities. We also demonstrate the value proposition of the assay by introducing MuSCs from aged animals and revealing multiple functional deficits tied to an aberrant quiescent state, which we show are rescued by wortmannin treatment, a quiescence-reinforcing strategy previously tested on “primed” MuSCs from young animals.^54^ These breakthroughs, together with the modularity of the assay components, miniaturized format, and validated semi-automated workflows to capture and process phenotypic data, offers an unprecedented opportunity to advance our understanding of MuSC quiescence and regulation in iterated designer niches.

In spite of a stress-induced response to tissue digestion and cell sorting^21^, our data demonstrate that the primary response of most MuSCs introduced to the biomimetic niche is immediate inactivation. A small proportion enter cell-cycle prior to inactivation and an even smaller subset directly differentiate and fuse with myotubes in the template. The MuSC population is increasingly regarded as encompassing a continuum of quiescence to activation^19^, and we believe our assay captures this continuum-influenced functional heterogeneity. For example, we expect those MuSCs closer to activation were inclined to differentiate and fuse, while those closer to a deeply quiescent state were resistant to activation cues. Additionally, Pax7^+^ donor cells at 7 DPE expressing CalcR and/or polarized niche markers represent a little over one-tenth of the population, hinting that a subset of more naive MuSCs are those recapitulating the more “advanced” hallmarks of quiescence we observed at 7 DPE. This is particularly intriguing taken with studies by others attributing a bonafide stem cell status to a similar proportion of MuSCs within the total population.^8,37,54,58^ Indeed, taken together with our observation that a subpopulation of engrafted donor MuSCs never enter cell cycle (**Figure 3**), we proport that the biomimetic niche maintains a “genuine-like” quiescent MuSC population alongside a more “primed-like” MuSC population, therein offering a tractable culture system with which to identify biochemical and biophysical regulators of these unique states.

Engrafted MuSCs showed population-level control over their response to the niche, which opens up enticing possibilities for studies of MuSC pool size regulation, and to uncover rules dictating niche repopulation. Of which increased differentiation was recently recognized as a quality control mechanism *in vivo*, similar to our own studies.^59^ As well, rules of niche occupancy also extoll limits on the number of transplanted MuSCs that can engraft into a recipient muscle.^60^ The data presented also underscores the importance of myotubes encased in a 3D matrix in allowing a persistent Pax7^+^ pool, and for determining the niche occupancy plateau point, despite a differentiation-inducing culture milieu and the absence of any other cell types. This is perhaps not surprising as many studies tout a role for myofibers in preserving or inducing quiescence, and in controlling MuSC pool size.^61–64^ A feedback mechanism between myofibers and MuSCs that is linked to nuclear content is suspected^64^; the likes of which could be interrogated in our system.

MuSCs *in situ* display long cytoplasmic projections that were initially described from electron microscopy analyses^65^, and more recently evaluated using tissue clearing and intravital imaging methodologies^12,47,48^. The elaborate MuSC morphologies arising in our cultures offer a new opportunity to explore the cellular and molecular mechanisms driving the acquisition of quiescent-like morphologies, but also the relevance of this phenotypic feature on MuSC behavior and fate. Indeed, long elaborated cytoplasmic projections have been associated with a deeper quiescent state^48^, and have been ruled out as a migratory apparatus^47^, favoring instead a role in ‘niche sensing’, though that remains to be determined.^47,48^ Quite surprisingly, we found that acquisition of quiescent-like morphologies and anatomical hallmarks was dependent on interactions between MuSCs and their immediate niche, occurring in the absence of other resident muscle cell types. The modular culture assay described herein enables iterative study design and independent molecular perturbations to the niche (myotubes) and the MuSCs to break open knowledge in this area. Indeed, leveraging high-content imaging and CellProfiler™ workflows for relating morphometric features to fate signatures, we offer proof of concept support for the use of morphological features as a non-invasive readout of MuSC quiescence status in our model, thereby facilitating future phenotypic screens.

To date, characterization of functional deficits of aged MuSC populations *in vitro* have focused on proliferation and colony formation as readouts^53,57,66–69^, and studies of other recognized deficiencies have been restricted to *in vivo* studies. In recent years, aged MuSC regenerative deficits have been linked to the notion that with age there is a progressive decrease in truly quiescent cells in favor of a greater number of cells in a pre-activated state.^20,55,56,70^ Consistent with functional consequences expected of a pre-activated state, we noted aberrant expansion activity at early culture timepoints from a subset of the aged MuSCs seeded within our assay, and a trend towards increased differentiation at later time-points, that were not observed in young MuSC cultures. Our studies also uncovered delayed inactivation kinetics and also in the acquisition of quiescent-like features. This suggests that a subset of aged MuSCs were unable to properly sense and respond to the pro-quiescent environment. Furthermore, the aged MuSC population was unable to maintain a steady-state pool size, which may imply that MuSC pool regulation is at least partly cell intrinsic and is dependent on both the activation state of MuSCs under steady-state conditions and exposure to activation-inducing cues. Interestingly, the myoblasts used to fabricate all of the muscle tissues for this study were derived from young mice, meaning that the aged MuSCs were exposed to a young niche. We cannot rule out the possibility that the young biomimetic muscle niche partially rescued aged MuSC function, as has been reported by others^71,72^. Indeed, we anticipate that aged MuSCs introduced to muscle tissues fabricated from myoblasts derived from aged donors and/or exposed to an aging systemic environment will induce further functional decline.

Finally, we found that inhibiting Akt signaling restores aged MuSC inactivation kinetics and population control, and partially rescues quiescent-like features. Our study extends prior work in showing that a strategy demonstrated to confer a genuine quiescent state onto young, activated MuSCs has a similar effect on aged MuSCs. We show that a decline in nuclear FoxOa3 levels is detected in MuSCs at an earlier age than previously thought, and that the nuclear FoxOa3 expression is corrected to youthlike levels by the wortmannin treatment. Wortmannin treated had only subtle influence on young MuSCs, which may reflect an absence stimulatory niche-derived ligands.^54^ However, we note that the DM culture media contains insulin, and the young and aged MuSCs were each cultured within young muscle tissues.

To conclude, herein we report a culture model capable of recapitulating aspects of quiescent MuSC biology, in youth and in age, that were previously not possible to study *in vitro*. By contrast to all other 3D culture systems where the cellular and ECM components are mixed together and introduced at the start of the experiment^27,29–32,73–75^, our method is modular. Amongst the merits of this distinction is the ability to introduce and evenly distribute new cellular components at any point in the assay. It is also feasible to genetically modify the MuSC and myoblast components in different ways, and maintain these distinctions, when MuSCs are introduced to the muscle tissue after myotubes have formed. These advantages, together with the simplicity of the approach, assay compatibility with existing semi-automated high content image acquisition and analysis tools, and high value features of MuSC biology captured by the system, offer a unique opportunity to expand MuSC fundamental knowledge and identify molecular targets to protect MuSC function as animals age.

## Materials and Methods

### Animal use protocols and ethics

All animal use protocols were reviewed and approved by the local Animal Care Committee (ACC) within the Division of Comparative Medicine (DCM) at the University of Toronto. All methods in this study were conducted as described in the approved animal use protocols (#20012838) and more broadly in accordance with the guidelines and regulations of the DCM ACC and the Canadian Council on Animal Care. 129-Tg(CAG-EYFP)7AC5Nagy/J (Actin-eYFP) mice^38^ were purchased from the Jackson Laboratory by the lab of Dr. Derek van der Kooy and shared with our group. Tg:Pax7-nEGFP (i.e. Pax7-nGFP) mice were a gift from Dr. Shahragim Tajbakhsh^76^, kindly transferred from the laboratory of Dr. Michael Rudnicki at the Ottawa Hospital Research Institute. Unless otherwise indicated, 8-12 weeks old mice were used for all experiments. A breeding pair of CB6-Tg(CAG-EGFP/CETN2)3-4Jgg/J (Centrin 2-eGFP) transgenic mice^77^ were kindly shared by Jeffrey Martens (University of Florida), and maintained by breeding for use in the aging studies. Young mice were between 4-5 months and aged mice between 24-26 months old.

### Magnetic-activated cell sorting (MACS) of primary mouse muscle stem cells

Primary mouse MuSCs were isolated from mouse hindlimb muscle using a modified method previously reported by our group.^33^ Briefly, ≈1 gram of muscle tissues was dissected from the hindlimb muscles of a humanely euthanized mouse and placed into a GentleMACS dissociation tube (Miltenyi Biotec, #130-096-334). 7 mL of DMEM (Gibco, #11995-073) with 630 U/mL Type 1A collagenase from clostridium histolyticum (Sigma, #C9891) was added to the tube, and the sample was physically dissociated using a GentleMACS dissociator (Miltenyi Biotec, #130-096-334) using the “skeletal muscle” setting. The tube was then placed on an orbital shaker in a 37 °C incubator for 1-hour. The digested tissue was triturated 10 times through a 10 mL pipette, after which an additional 440 U of Type 1A collagenase was added along with Dispase II (Life Technologies, #17105041) and DNAse I (Bio Basic, #9003-98-9) at a final concentration of 0.04 U/mL and 100 µg/mL, respectively. The tube was again placed on an orbital shaker in a 37 °C incubator for 1-hour. The sample was then slowly passed through a 20 G needle 15 times and then resuspended in 7 mL of FACS buffer (**Supplementary Table 1**). The solution was passed through a 70 µm cell strainer (Miltenyi Biotec, #130-098-462) followed by a 40 µm cell strainer (Corning, #352340). The filtered mixture was then centrifuged at 400 g for 15 minutes and the supernatant aspirated. The pellet was resuspended in 1 mL of 1X red blood cell (RBC) lysis buffer (**Supplementary Table 1**) and then incubated at room temperature (RT) for 8 minutes. 9 mL of FACS buffer was added to the tube and the mixture was centrifuged at 400 g for 15 minutes followed by supernatant aspiration.

The cell pellet was then incubated in a 4 °C fridge with rocking for 15 minutes in 100 µL of MACS buffer and 25 µL of lineage depletion microbeads from the Satellite Cell Isolation Kit (Miltenyi Biotec, #130-104-268) according to the manufacturer’s instructions. Another 375 µL of MACS buffer was then added, and the lineage positive cells depleted by flowing the solution, by gravity, through an LS column in a magnetic field (Miltenyi Biotec, #130-042-401) (**Supplementary Table 1**). The resulting flow through was collected, corrected to 5 mL and then centrifuged at 400 g for 5 minutes. The pellet was then subjected to a second round of lineage depletion using a fresh LS column in a magnetic field. The flow through was corrected to 5 mL, centrifuged, followed by supernatant aspiration, and then the cell pellet was resuspended in 100 µL of MACS buffer and 25 µL of anti-integrin α-7 microbeads (Miltenyi Biotec, #130-104-261) for incubation at 4 °C for 15 minutes. 375 µL MACS buffer was added, and the integrin α-7^+^ was enriched by running the solution through a third LS column in a magnetic field. In this instance, the flow through was discarded, the column was removed from the magnetic field and then flushed with 5 mL of MACS buffer which was collected in a 15 mL conical tube. The tube was spun to generate a cell pellet enriched for integrin α-7^+^ MuSCs. To establish and validate the protocol, which differs from the manufacturers protocol by the introduction of extra lineage depletion steps, α-7^+^ MuSCs were isolated from Pax7-nGFP transgenic mice. In these experiments the cell pellet was resuspended in 0.5 mL FACS buffer and incubated with DRAQ5 for 15 min at RT. After 3 × 5 min FACS buffer washes and centrifuge spins, the pellet was resuspended in 0.5 mL of FACS buffer and propidium iodide (PI) was added to the tube. The resuspended cells were then evaluated using the Accuri C6 Flow Cytometer (BD Biosciences) whereby we collected 30,000 events. The DRAQ5^+^Pax-nGFP^+^PI^-^ cell population was quantified from the flow cytometric data using FlowJo™ V10 software.

### Primary mouse myoblast line derivation and maintenance

Prior myoblast cell lines were derived from freshly MACS enriched integrin α-7^+^ MuSC populations. 1-day before cell plating, culture dishes were coated at 4 °C overnight with collagen I at a 1:8 concentration diluted in ddH_2_O (Gibco, #A10483-01). The next day, excess collagen I solution was removed, and the dish culture surfaces were dried at RT for 15-20 min followed by a PBS wash prior to use. Immediately after MACS isolation, lineage depleted integrin α-7^+^ enriched MuSCs were resuspended in SAT10 media (**Supplementary Table 1**) and plated into collagen-coated dishes. A full media change was performed 48 hours after plating with half media changes every 2 days thereafter. Cells were grown to 70 % confluency and passaged at least 5 times to produce a primary mouse myoblast line, and then used from passage 5 – 9 for experiments.

### Murine myotube template fabrication and MuSC seeding

One day prior to seeding myotube templates, black 96-well clear bottom plates (PerkinElmer, #6055300) were coated with 5 % pluronic acid (Sigma-Aldrich, #P2443) and incubated overnight at 4 °C. The next day, excess pluronic solution was removed, and plates were left at RT for 15 – 20 min to dry well surfaces. Cellulose paper (MiniMinit) was cut into 5 mm discs using a biopsy punch (Integra, #MLT3335), autoclaved, and then placed into pluronic acid coated wells of the 96-well plate. A stock thrombin solution (100 U/mL, Sigma-Aldrich, #T6884) was then diluted to 0.8 U/mL in PBS, and then 4 µL was diffused into the paper discs and left to dry at RT. Meanwhile, a 10 mg/mL fibrinogen solution was made by dissolving lyophilized fibrinogen (Sigma-Aldrich, #F8630) in a 0.9 % wt/vol solution of NaCl (Sigma-Aldrich, #S5886) and then filtered through a 0.22 µm syringe filter (Sarstedt, #83.1826.001). Primary myoblasts were then trypsinized, counted using a hemacytometer, and then resuspended in an ECM-mimicking slurry comprised of 40 % DMEM, 40 % Fibrinogen, and 20 % Geltrex™ (ThermoFisher, #A1413202) at a concentration of 25,000 cells per 4 µL. The cell / extracellular matrix solution was then diffused into dry thrombin-containing paper discs and left to gel at 37 °C for 5 min. 200 µL growth media (GM, **Supplementary Table 1**) was introduced to each hydrogel containing culture well and plates were returned to a cell culture incubator (37 °C, 5 % CO_2_) for 2 days (Day -2 to 0). On Day 0 of differentiation, a full media change was conducted to transition cultures to differentiation media (DM, **Supplementary Table 1**). Half media changes with DM were performed every other day from thereafter.

Unless otherwise indicated, on Day 5 of myotube template culture integrin α-7^+^ MuSCs were prospectively isolated and resuspended in SAT10 media replete of FGF2. Myotube templates were carefully removed from the 96-well plate using tweezers and placed in an ethanol-sterilized plastic container containing long strips of polydimethylsiloxane (PDMS) sitting on top of a moist paper towel. Quickly, 4 µL of the resuspended MuSC solution containing the desired number of MuSCs was placed onto each tissue and evenly spread over the tissue surface using a cell-spreader. The plastic container was then sealed with a tight fitting lid and placed in the 37 °C incubator for 1 hour before putting the tissues back into their wells using tweezers. For aged MuSC-related studies, minced and cryopreserved hindlimb muscle from young or litter-matched aged Centrin 2-eGFP mice^77^ were thawed and underwent the MACS protocol detailed above. The MuSCs were then resuspended in SAT10 media replete of FGF2 but with added wortmannin (10 µM, Sigma-Aldrich, #W1628) or a dimethyl sulfoxide (DMSO) control (Sigma-Aldrich, #D8418). 4 µL of the resuspended MuSCs containing ≈500 cells were subsequently seeded onto individual tissues. After 1 hour, tissues were put back into their wells. The wortmannin (or DMSO) was then added to the culture media (also at 10 µM) and refreshed every other day during media changes.

### Tissue fixation and immunolabelling

At the indicated tissue endpoints, samples were quickly washed 3x with PBS before fixation with 100 µL of 4 % paraformaldehyde (PFA, Fisher scientific, #50980494) for 12 min at RT. After 3 × 10 min washes with cold PBS (4 °C), blocking and permeabilization was performed using 100 µL of blocking solution (**Supplementary Table 1**) for 30 min at RT. Afterwards, primary antibodies were diluted in blocking solution as indicated in **Supplementary Table 2** and 50 µL was added to each tissue and incubated overnight at 4 °C. After 3 × 10min washes with cold PBS, tissues were incubated for 45 min at RT in 50 µL of secondary antibodies and molecular probes diluted in blocking solution (see **Supplementary Table 2**), followed by 3 × 10 min washes with cold PBS. A limitation of the cellulose papers is that they cast autofluorescence in the blue channel, which can give off intense background noise. Therefore, for nuclei detection, DAPI was sometimes used as the signal intensity was generally high enough to allow thresholding of paper fibers out of confocal images. Batch to batch differences in DAPI, or in cases when tissues become dry during staining, can result in DAPI images where the cellulose fibers are visualized, although even in these cases the nuclei can still be clearly discerned.

### Image acquisition

Confocal imaging was performed using the Perkin-Elmer Operetta CLS High-Content Analysis System and the associated Harmony® software. Prior inserting the 96-well plate into the Operetta, the PBS was removed from the wells of the plate to prevent tissues from shifting during imaging, and they were carefully positioned in the middle of the wells using tweezers. For stitched pictures, images were collected using the 10X air objective (Two Peak autofocus, NA 1.0 and Binning of 1). For MuSC analysis, images were collected using the 20X and 40X water immersion objectives (Two Peak autofocus, NA 1.0 and 1.1, and Binning of 1). All images were exported off the Harmony® software in their raw form. Subsequent stitching, max projections, etc was performed using the ImageJ-BIOP Operetta Import Plugin available on c4science.^78^ For imaging of MuSC niche markers, the Olympus FV-1000 confocal microscope and Olympus FluoView V4.2b imaging software was used along with a 40X silicone immersion objective (NA 1.25; Olympus, #UPLSAPO40XS).

### Bio-image analysis

For SAA coverage, stitched images were used along with a previously published ImageJ macro.^79^ The SAA signal was put in red, the threshold set to 0-45 and the tissue outline selected using the oval tool. For fusion index, cell counting, cell morphology, YFP/GFP coverage and mean nuclear intensity, the CellProfiler™ software was utilized. CellProfiler™ version 4.2.1^49^ was downloaded from source website (www.cellprofiler.org) and installed on a PC (Intel Core i9-11900 @ 2.5GHz, 64.0 GB RAM, and 64-bit Windows 11 operating system). Analysis pipelines were created for each of the above-mentioned metrics. *Fusion index*: 9 × 20X max projected images were taken per tissue. The channels were split, the fiber and nuclei signal individually identified and overlayed to calculate the percentage of nuclei in fibers. *Cell counting and morphology*: 25 × 20X max projected images were taken per tissue. The channels were split, mononucleated DAPI^+^YFP^+^Pax7^+^ (or Caveolin-1^+^) objects extracted using the IdentifyPrimaryObjects module and counted. For object segmentation, the global minimum cross-entropy thresholding method^80^ was selected. Pixel intensity and object shape were used as metrics to distinguish and segment clumped objects. Morphology measurements of the identified cellular objects were recorded using the MeasureObjectSizeShape module. For the proportion of c-FOS^+^, Ki67^+^, MyoD^+^ and/or CalcR^+^ cells, this fourth channel was overlayed over the identified objects and divided. *YFP/GFP coverage*: 25 × 20X max projected images were taken per tissue. The channels were split, the YFP/GFP signal identified, and coverage calculated using the MeasureAreaOccupied function. *Mean nuclear intensity*: 104 × 40X max projected images were taken per well. The channels were split, the nuclei, cell and cytoplasm identified as primary, secondar and tertiary objects, and the intensity of the FoxO3a signal within the nuclei calculated using the MeasureObjectIntensity function.

### MTS assay

To quantify the metabolic activity of myotube templates, the MTS assay was used (abcam, #ab197010). First, 200 µL of fresh DM was added to each tissue. Then, 20 µL of the MTS tetrazolium compound was added to each well and incubated for 2 hours at 37 °C. The media was vigorously mixed with a pipette every 30 min to ensure maximal diffusion of the formazan dye product. The entire culture media from each tissue was then pipetted into a clear 96-well plate (Sarstedt, #83.3924) and the OD at 490 nm quantified with a spectrophotometer (Tecan, Infinite M200 Pro). The assay was performed on different tissues on different days of culture, each with their own “media + MTS” negative control, which was subtracted as background from all OD values.

### EdU assay experiments

EdU experiments were performed using the Invitrogen Click-iT™ Plus EdU Alexa Fluor™ 555 Imaging Kit (#C10638). EdU was added to the culture media on Day 5 after MuSC engraftment and refreshed every 24 hours until 7 DPE. After tissue fixation and blocking, EdU labelling was done according to the product protocol apart from a 20 min incubation instead of 30 min. Subsequent immunolabelling was done as described above. The CellProfiler™ pipeline was then implemented to identify mononucleated DAPI^+^YFP^+^Ki67^-^ objects and then overlayed with the EdU channel to quantify the proportion of EdU positive cells.

### Barium chloride tissue injury

On Day 12 of differentiation (7 DPE), the culture media was removed, and tissues were incubated with either PSS (**Supplementary Table 1**) or a 2.4 % wt/v BaCl_2_ solution diluted in PSS for a period of 4 hours (protocol adapted from previously published literature^81^). Tissues were then washed 3 × 5 min with warm wash media (**Supplementary Table 1**) and then returned to fresh DM for 2 more days before fixation.

### 2D culture experiments

For 2D myotube culture experiments, microwells were first coated with a 5 % v/v Geltrex™/DMEM solution for 1 hour at 37 °C. After drying, 25,000 primary myoblasts were added per well in 200 μL of GM, which was then switched to DM after 2 days. On day 5 of differentiation, 500 MuSCs were engrafted onto 2D myotubes in a 4 μL volume.

### Statistical analysis

Statistical analysis was performed using the GraphPad Prism 9 software. Most experiments were performed with 3 technical tissue replicates per experimental group and repeated on 3 independent occasions (i.e., n=9 technical replicates across N=3 biological replicates). Please refer to **Supplementary Table 3** for a specific breakdown of replicates per experiment. All error bars show standard error of the mean (SEM). Significance was defined as p ≤ 0.05

## Acknowledgements

This project was funded by a CIHR CGS-D scholarship to E.J., an Ontario Graduate Scholarship to E.J., a Cecil Yip Doctoral Award to E.J., a Medicine by Design Canada First Research Excellence Fund grant to P.M.G. (MbDC2-2019-02), and a Canada Research Chair in Endogenous Repair to P.M.G.

## Author contributions

E.J. and PMG conceived of the project. E.J. and Y.K. designed and performed research, analyzed data, and prepared figures. P.M.G. supervised the research. All authors contributed to data interpretation. E.J., Y.K., and P.M.G. wrote the manuscript. All authors reviewed and approved the manuscript.

