## Supplementary Information for "Rescue of aged muscle stem cell intrinsic quiescence defects by AKT inhibition revealed with a 3D biomimetic culture assay"

**This PDF file includes:**

Supplementary Figures 1-13

Supplementary Tables 1-3

### Supplementary Figure 1

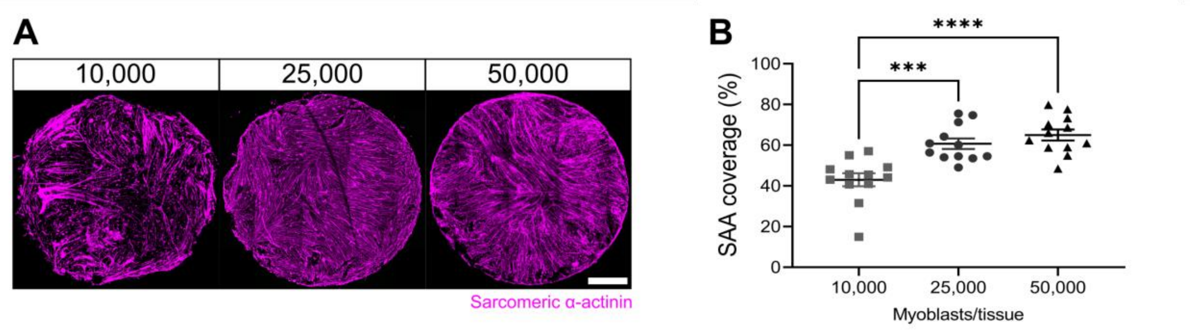


**Supplementary Figure 1. Optimization of cell seeding density. (A)** Representative confocal stitched images of myotube templates labelled for sarcomeric α-actinin (SAA; magenta) at day 5 using 10,000; 25,000; or 50,000 myoblasts per tissue. Scale bar, 1 mm. **(B)** Quantification of SAA area coverage of myotube templates at day 5 of differentiation using 10,000; 25,000; or 50,000 cells. n=12 across N=3 independent biological replicates. Graph displays technical replicates with mean ± s.e.m.; one-way ANOVA with Tukey post-test, *** p=0.0003 **** p<0.0001.

### Supplementary Figure 2

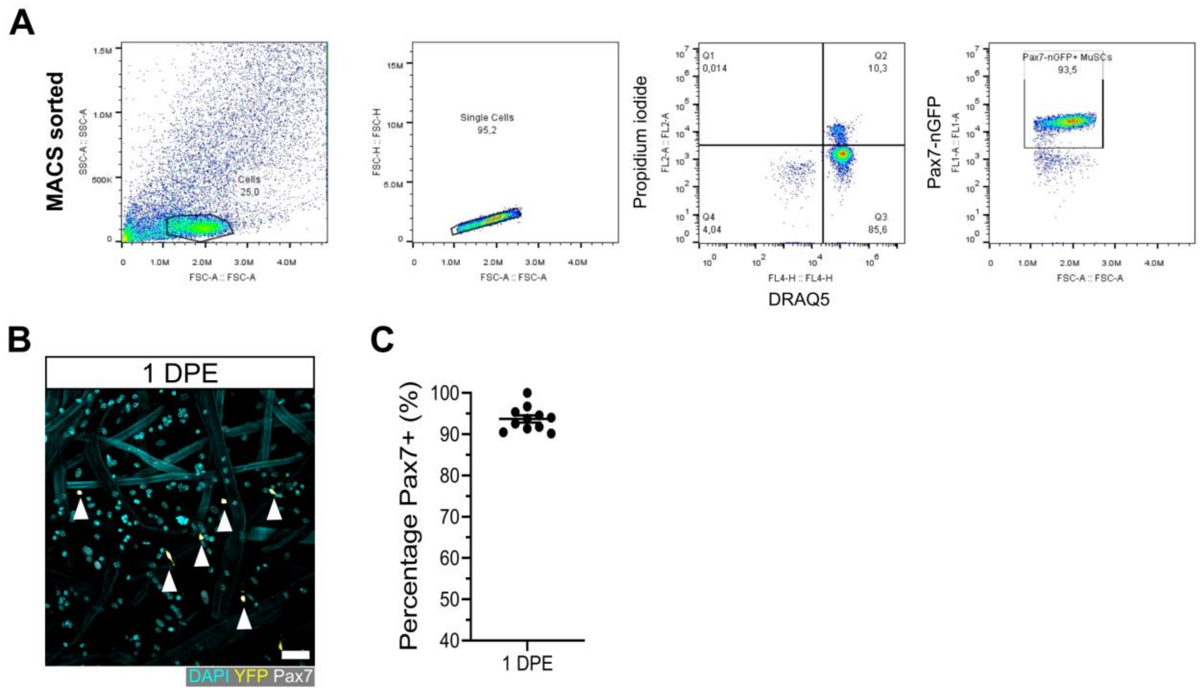


**Supplementary Figure 2. Population purity in MACS isolated MuSCs. (A)** Flow cytometry gating of MuSCs enriched from the hindlimb muscles of a Pax7-nGFP mouse using the MACS-based workflow. **(B)** Representative confocal image of MuSCs (DAPI: cyan, YFP: yellow, Pax7: white, white arrows) at 1 DPE onto myotube templates. Scale bar, 50µm. **(C)** Quantification of the percentage Pax7$^{+}$ cells in the DAPI$^{+}$YFP$^{+}$ population at 1 DPE after engraftment with CAG-EYFP reporter MuSCs. n=11 across N=3 independent biological replicates. Graph displays mean ± s.e.m. of technical replicates.

### Supplementary Figure 3

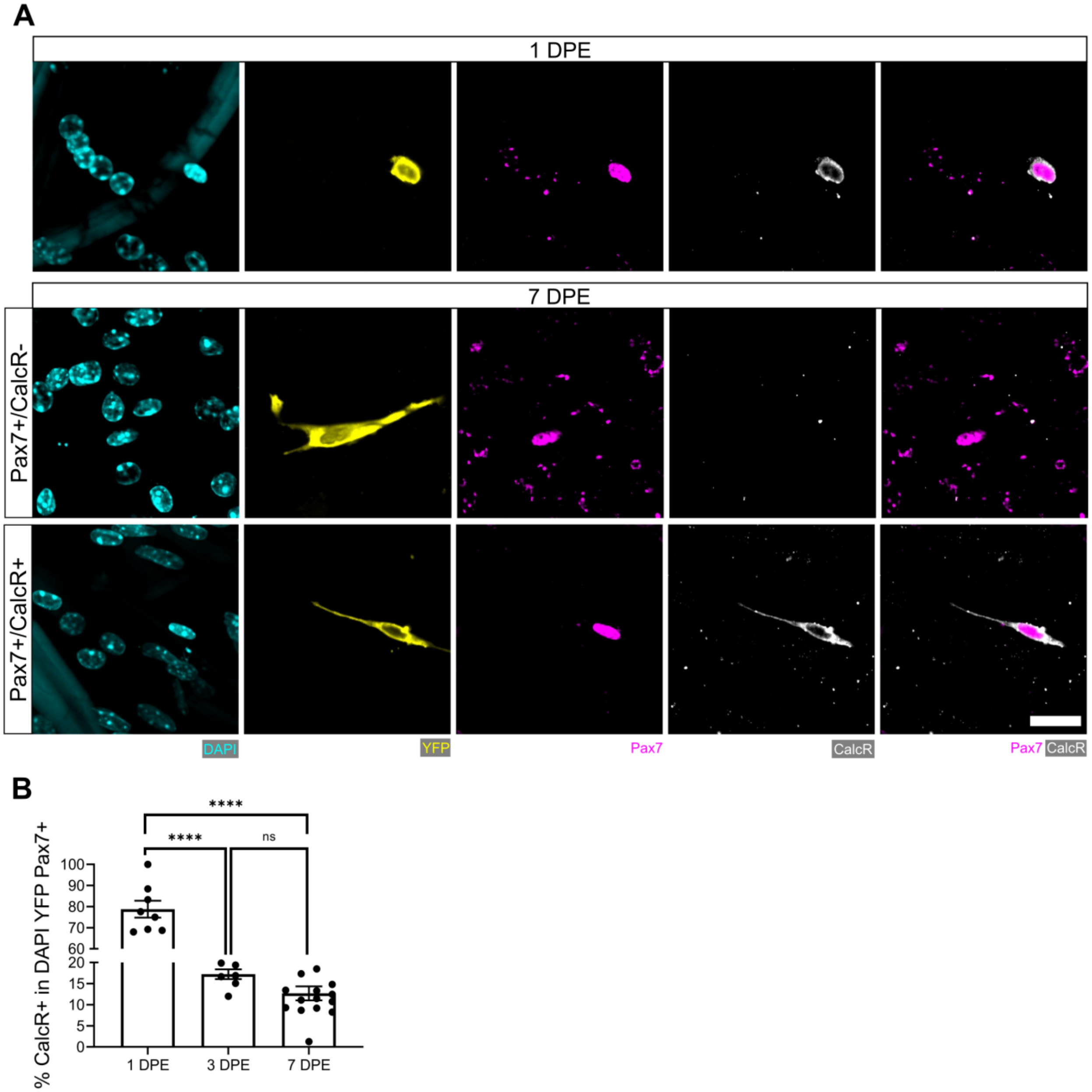


**Supplementary Figure 3. Persistent CalcR$^+$ population amongst Pax7$^+$ donor cell population at 7 DPE. (A)** Representative confocal images of a Pax7$^+$ (magenta) donor cells (yellow) at 1 DPE (top) and 7 DPE (middle and bottom) immunostained for calcitonin receptor (CalcR: white) and counterstained with DAPI (cyan). Scale bar, 20 µm. **(B)** Bar graph showing the percentage of CalcR$^+$ cells in the DAPI$^+$YFP$^+$Pax7$^+$ mononucleated cell population at 1, 3 and 7 DPE. n=7-15 across N=3-5 independent biological replicates. Graph displays mean ± s.e.m. of the individual technical replicates; one-way ANOVA with Tukey's post-test, **** $p<0.0001$.

### Supplementary Figure 4

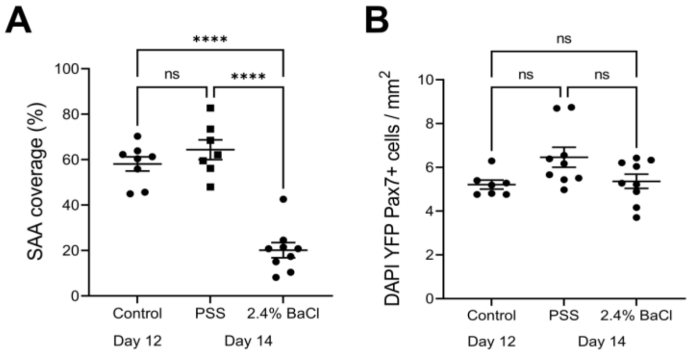


**Supplementary Figure 4. Characterization of barium chloride-induced injury. (A)** Quantification of SAA coverage on day 12 of differentiation, and 2 days (day 14) after a physiological salt solution (PSS; carrier control) or 2.4 % $BaCl_2$ exposure. n=7-9 across N=3 independent biological replicates. Graph displays mean ± s.e.m. with individual technical replicates; one-way ANOVA with Tukey post-test, **** $p<0.0001$. **(B)** Quantification of $DAPI^+YFP^+Pax7^+$ mononucleated cell density on day 12 of differentiation, and 2 days (day 14) after a PSS or 2.4 % $BaCl_2$ exposure. n=6-9 across N=2-3 independent biological replicates. Graph displays mean ± s.e.m. with individual technical replicates; one-way ANOVA, non-significant (ns).

### Supplementary Figure 5

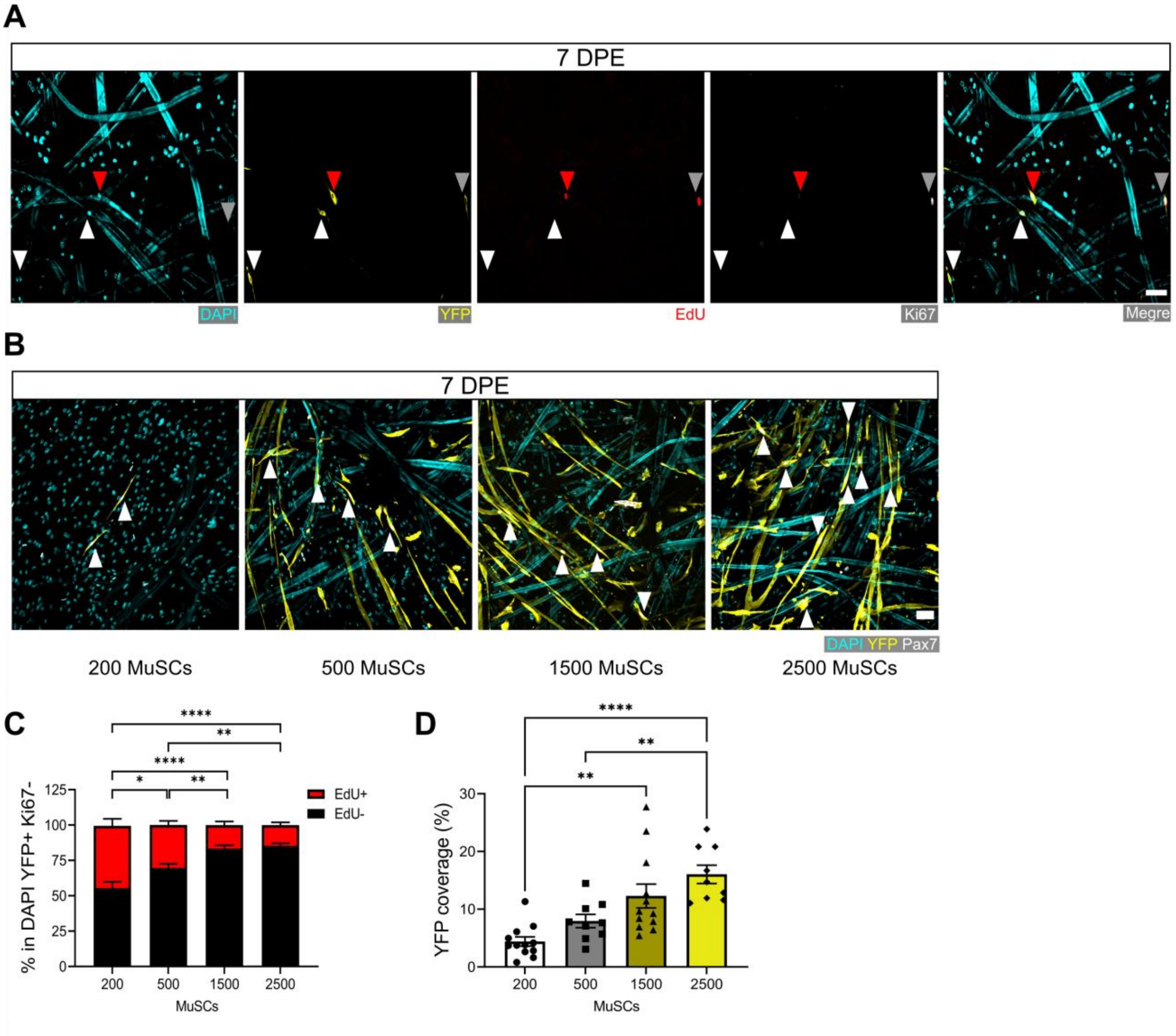


**Supplementary Figure 5. Regulation of MuSC pool size in myotube template cultures. (A)** Representative confocal image of YFP$^{+}$ (yellow) mononucleate donor cells at 7 DPE co-labeled for EdU (red), Ki67 (white) and nuclei (DAPI: cyan), to visualize EdU$^{+}$Ki67$^{-}$ (red arrow), EdU$^{+}$Ki67$^{+}$ (white, grey arrow), and EdU$^{-}$Ki67$^{-}$ (white arrows) populations. Scale bar, 50µm. **(B)** Representative confocal images of tissues at 7 DPE initially seeded with 200, 500, 1500 or 2500 MuSCs (DAPI: cyan, YFP: yellow, Pax7: white, white arrows). Scale bar, 50 µm. **(C)** Proportion of EdU+/- cells at 7 DPE in the DAPI$^{+}$YFP$^{+}$Ki67$^{-}$ population across different starting MuSC engraftment numbers (200, 500, 1500 or 2500). n=10-16 across N=3-5 independent biological replicates. Graph displays mean ± s.e.m. for EdU$^{+}$ and EdU$^{-}$; one-way ANOVA with Tukey's post-test comparing the EdU$^{-}$ proportions of each condition, * p=0.0102 ** p=0.0063, 0.0026 **** p<0.0001. **(D)** Quantification of tissue area covered by YFP signal at 7 DPE when engrafted with 200, 500, 1500 or 2500 MuSCs. n=9-12 across N=3-4 independent biological replicates. Graph displays mean ± s.e.m. with individual technical replicates; one-way ANOVA with Tukey post-test, ** p=0.0019, 0.0066 **** p<0.0001.

### Supplementary Figure 6

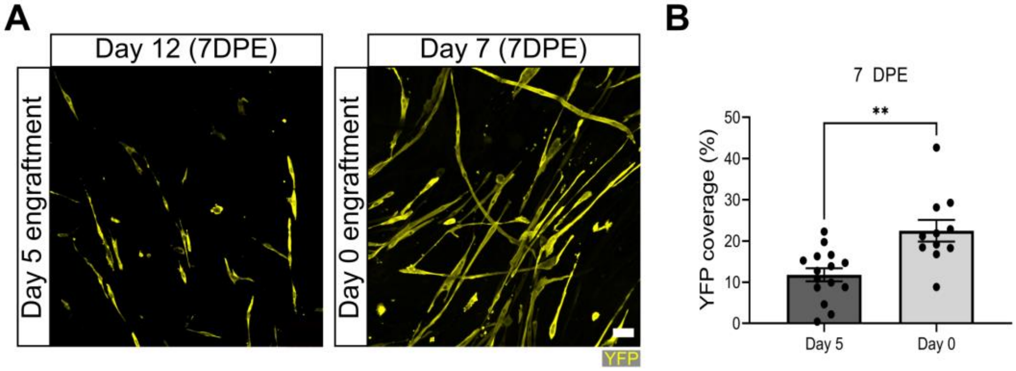


**Supplementary Figure 6. Increased YFP coverage when MuSCs engrafted on Day 0 of myotube template differentiation. (A)** Representative confocal images of tissues engrafted with 500 MuSCs at day 0 or day 5 of myotube template differentiation, fixed at 7 DPE (day 7 and 12), and immunolabelled for YFP (yellow). Scale bar, 50 µm. **(B)** Quantification of percentage of tissue area covered by YFP signal at 7 DPE when 500 MuSCs are engrafted on day 0 versus day 5. n=11, 15 from N=4, 5 independent biological replicates. Graph displays mean ± s.e.m. with individual technical replicates; unpaired t-test, ** p=0.0012.

### Supplementary Figure 7

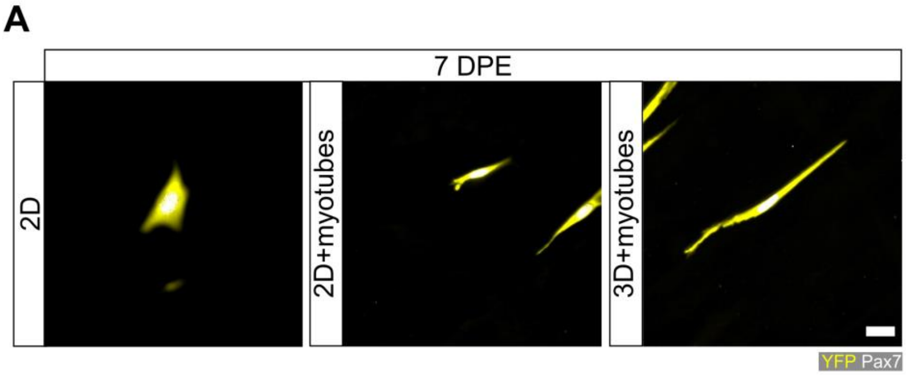


**Supplementary Figure 7. Pax7$^{+}$ donor cell morphologies in 2D and in 3D. (A)** Representative confocal images of mononucleated donor cells (YFP:yellow; Pax7:white) after 7 days in a 2D collagen I-coated petri dish (left), in 2D culture with a myotube monolayer and a Geltrex™ undercoating (middle), or engrafted onto myotube templates at Day 5 of differentiation (right). Scale bar, 20 µm.

### Supplementary Figure 8

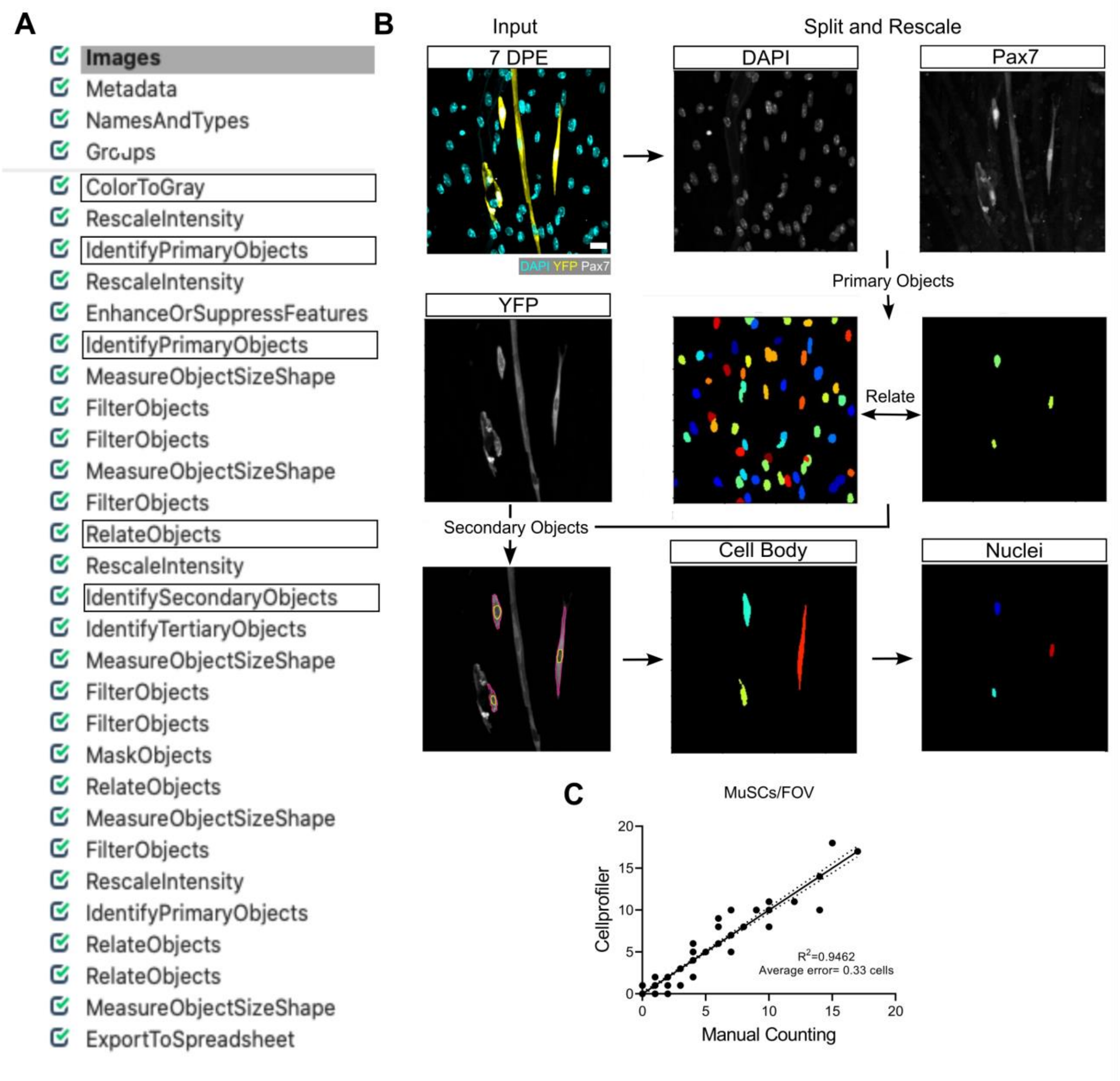


**Supplementary Figure 8. CellProfiler™ pipeline for MuSC identification and characterization. (A)** Representative image of the pipeline loaded in CellProfiler™. **(B)** Representative confocal image of mononucleated (DAPI:cyan) donor (YFP:yellow) MuSCs (Pax7:white) at 7 DPE (white arrowheads) along with a general schematic of the CellProfiler™ workflow (see Methods for more detail). Scale bar, 20 µm. **(C)** Dot-plot graph showing the correlation between Manual and CellProfiler™ counting of MuSCs in individual images (FOV). n=125 images across N=3 independent biological replicates, graph displays individual technical replicates (images) with a simple linear regression analysis (solid line) and 95% confidence intervals (dotted lines). Remark: many of the 125 datapoints in (C) overlap.

**Supplementary Figure 9**

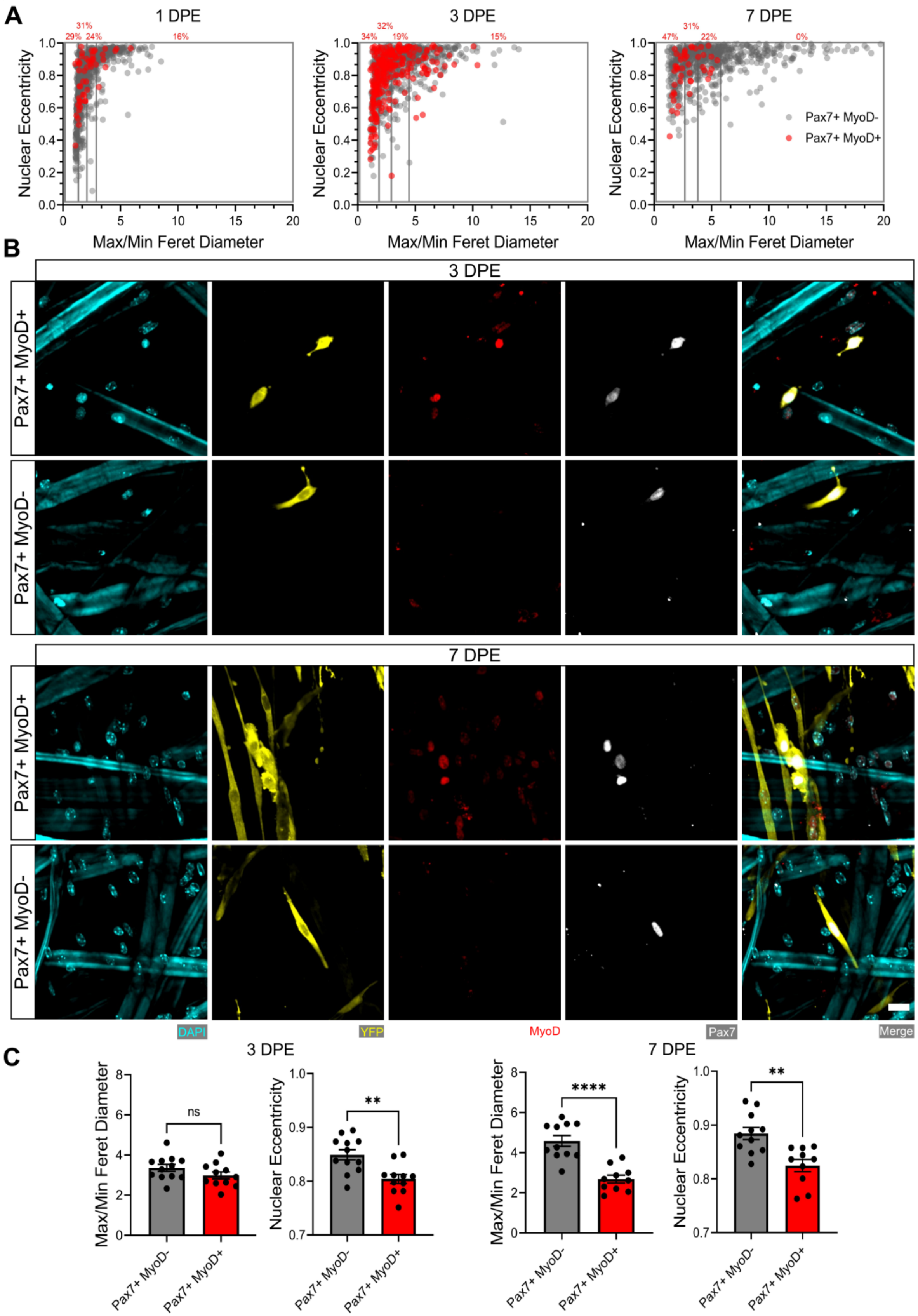
A
1 DPE
3 DPE
7 DPE
Nuclear Eccentricity
Max/Min Feret Diameter
31%
29%
24%
16%
32%
34%
19%
15%
47%
22%
0%
Pax7+ MyoD-
Pax7+ MyoD+
B
3 DPE
7 DPE
Pax7+ MyoD+
Pax7+ MyoD-
DAPI
YFP
MyoD
Pax7
Merge
C
3 DPE
7 DPE
ns
**
****
Max/Min Feret Diameter
Nuclear Eccentricity

**Supplementary Figure 9. Morphological characterization of Pax7$^{+}$MyoD$^{-}$ and Pax7$^{+}$MyoD$^{+}$ MuSCs. (A)** Dot-plot graphs showing individual Pax7$^{+}$ donor cells and their associated max/min feret diameter ratio and nuclear eccentricity at 1 (left), 3 (middle), and 7 DPE (right) with Pax7$^{+}$MyoD$^{+}$ cells shown in red and Pax7$^{+}$MyoD$^{-}$ cells in grey. Quartiles are indicated by grey boxes and the percentage of Pax7$^{+}$MyoD$^{+}$ cells within each quartile is written above in red. n=916, 980 and 737 across N=3-4 biological replicates also analyzed in Figure 5C. **(B)** Representative confocal images of donor MuSCs (DAPI:cyan; YFP:yellow; Pax7:white) +/- MyoD (red) at 3 (top) and 7 DPE (bottom). Scale bar, 20 µm. **(C)** Bar graphs showing the average max/min feret diameter ratio and nuclear eccentricity between Pax7$^{+}$MyoD$^{-}$ (dark grey) and Pax7$^{+}$MyoD$^{+}$ cells (red) at 3 (left) and 7 DPE (right). n=10-12 across N=3-4 independent technical replicates. Graphs display mean ± s.e.m. of the individual technical replicate tissues from panel A; unpaired t-test, ** p=0.0021, 0.0015 **** p<0.0001.

### Supplementary Figure 10

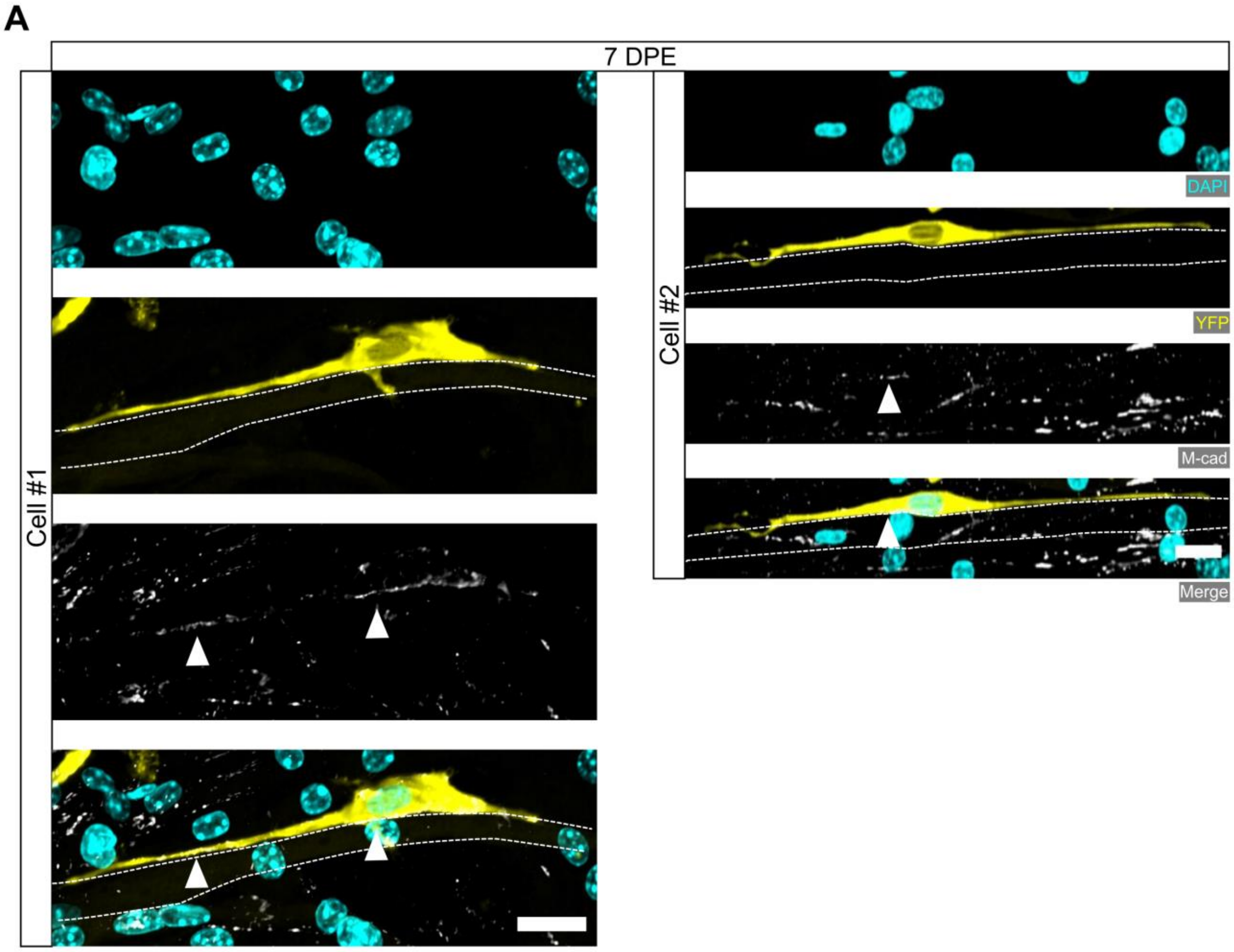


**Supplementary Figure 10. Polarized localization of M-cadherin in mononuclear donor cells at 7 DPE. (A)** Representative confocal images of two (Cell #1, left; Cell #2, right) mononucleated donor cells (DAPI: cyan; YFP: yellow) with M-cadherin (M-cad: white, white arrows) labeling restricted to the apical side. Cell #1 is associated with a dimly YFP$^{+}$ myotube (white dotted lines). Scale bars, 20 µm.

### Supplementary Figure 11

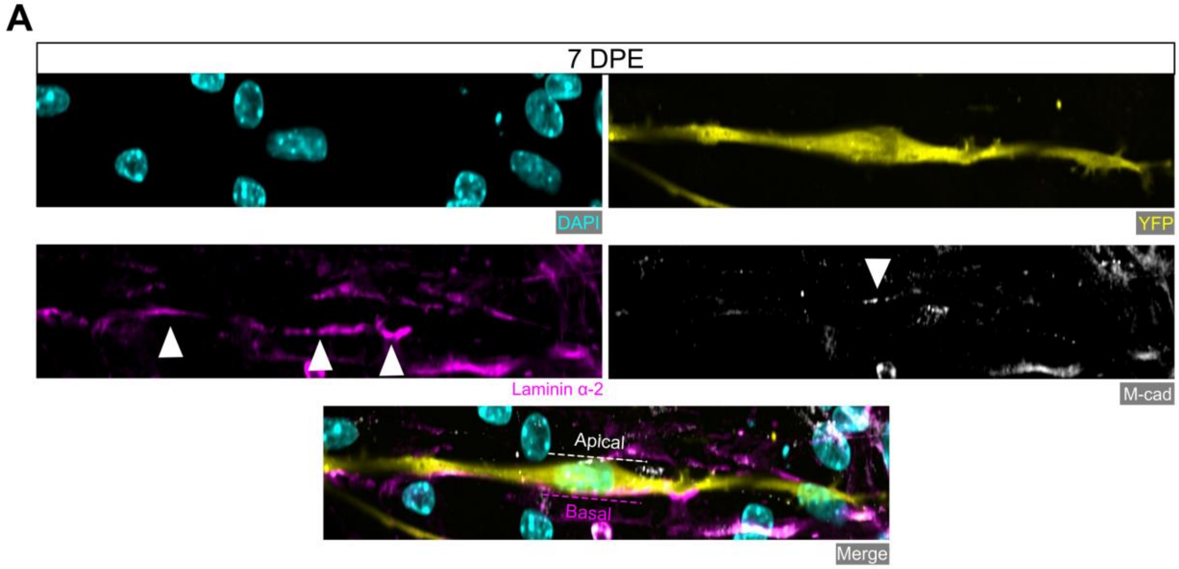


**Supplementary Figure 11. Donor cell polarization of niche markers at 7 DPE. (A)** Representative confocal image of a mononucleated donor cell (DAPI: cyan; YFP: yellow) with M-cadherin (white) immunolabelling restricted to the apical side and laminin α-2 (magenta) localized to the basal side. Scale bar, 20 µm.

### Supplementary Figure 12

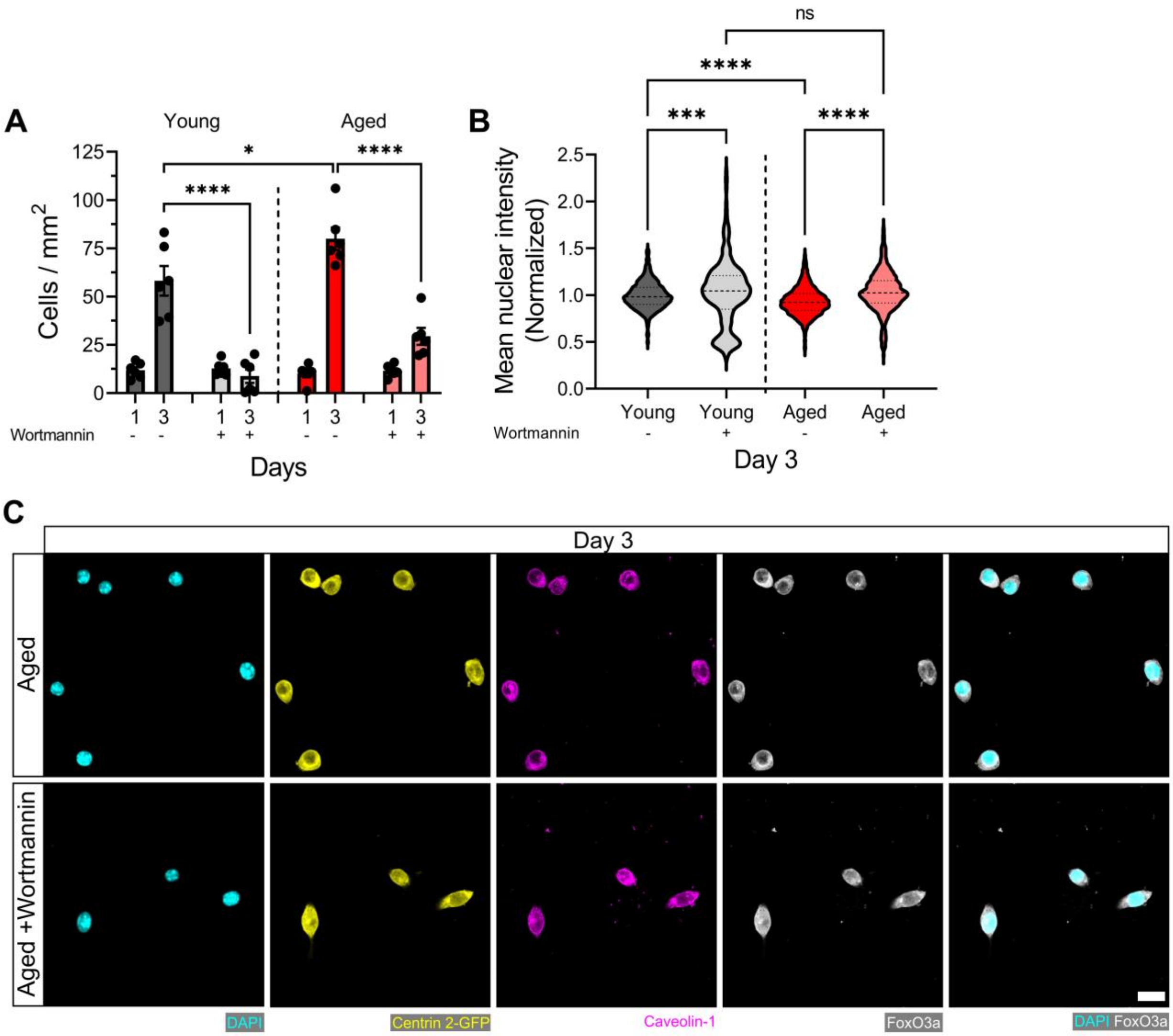


**Supplementary Figure 12. Wortmannin treatment blunts cell proliferation and increases FoxO3a nuclear localization in young and aged MuSCs cultured *in vitro*. (A)** Bar graph showing the mean number of MuSCs per mm$^2$ of 2D microwells on Day 1 and 3 across experimental conditions (young: dark grey; young + wortmannin: light grey; aged: red; aged + wortmannin: light red). n=6 across N=2 independent biological replicates. Graph displays mean ± s.e.m. with the individual technical replicates; one-way ANOVA with Tukey's post-test comparing each experimental group at the 3 DPE timepoint, * p=0.0353 **** p<0.0001. **(B)** Violin plot showing the mean nuclear fluorescent intensity of FoxO3a in MuSCs cultured in 2D microwells on Day 3 across experimental conditions (young: dark grey; young + wortmannin: light grey; aged: red; aged + wortmannin: light red). n=2716, 565, 4437, 1897 across N=2 independent biological replicates. Graph displays mean with first and third quartiles. Data was normalized to the average intensity of the young condition and outliers identified using the ROUT method (with Q=1%) were removed; one-way ANOVA with Šidák's post-test comparing pre-selected conditions, **** p=0.0002 **** p<0.0001.

### Supplementary Figure 13

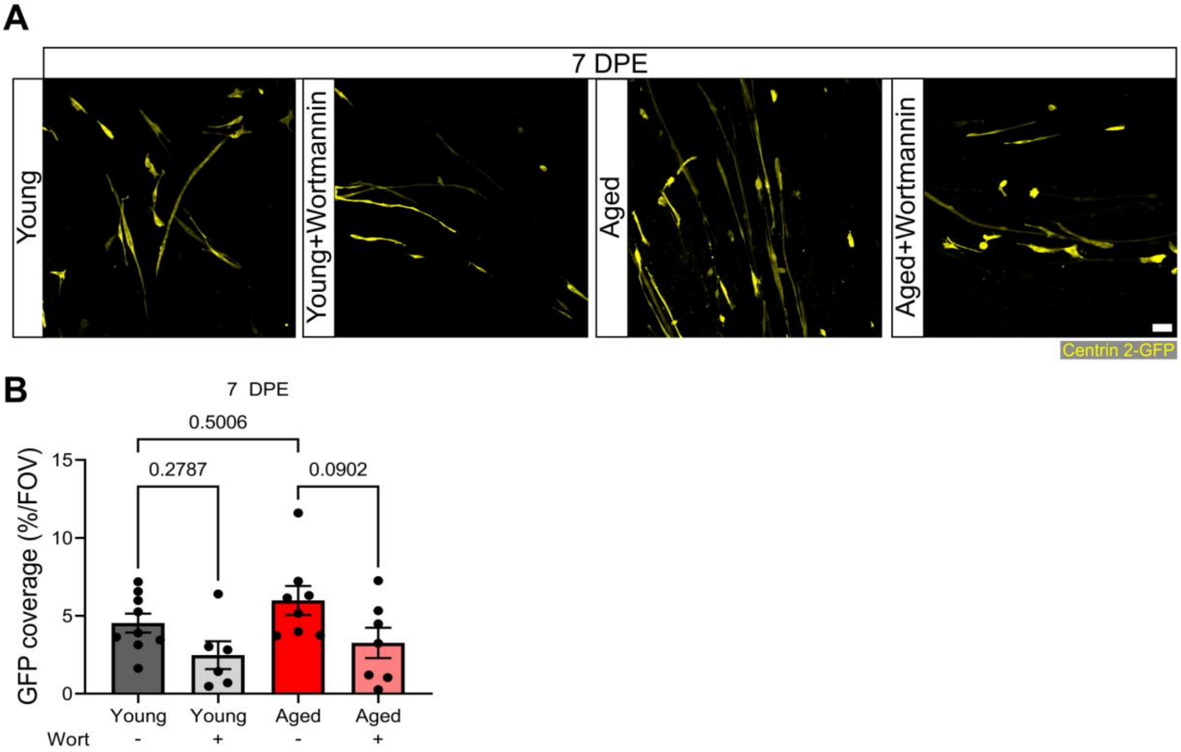


**Supplementary Figure 13. Wortmannin treatment diminishes GFP coverage at 7 DPE. (A)** Representative confocal images of the Centrin 2-GFP area coverage (yellow) at 7 DPE from young and aged engrafted MuSCs treated with DMSO control or wortmannin. Scale bar, 50 µm. **(B)** Bar graph showing the average GFP coverage (from the Centrin 2-GFP reporter) per field of view at 7 DPE across experimental conditions (young: dark grey; young + wortmannin: light grey; aged: red; aged + wortmannin: light red). n=6-9 across N=2-3 independent biological replicates. Graph displays mean ± s.e.m. with the individual technical replicates; one-way ANOVA with Tukey's post-test (comparisons not shown are not significant).

**Supplementary Table 1**

| Media | Composition |
|---|---|
| FACS Buffer | PBS, 2.5 % Goat serum (Gibco, #16210072), 2 mM EDTA (Sigma-Aldrich, #E5134) |
| RBC Lysis Buffer | $ddH_2O$, 0.155 M $NH_4Cl$ (Sigma-Aldrich, #A9434), 0.01 M $KHCO_3$ (Sigma-Aldrich, #237205), 0.1 mM EDTA |
| MACS Buffer | PBS, 0.5% Bovine serum albumin (BioShop, #9048-46-8), 2 mM EDTA |
| SAT10 | DMEM/F12 (Gibco, #11320-033), 1% Penicillin-Streptomycin (Gibco, #15140-122), 20% Fetal bovine serum (Gibco, 12483-020), 10% Horse serum (Gibco, #16050-122), 1% Glutamax (Gibco, #35050-061), 1% Insulin-Transferrin-Selenium (Gibco, #41400-045), 1% Non-essential Amino acids (Gibco, #11140-050), 1% Sodium Pyruvate (Gibco, #11360-070), 50µM b-mercaptoethanol (Gibco, #21985-023), 5ng/mL bFGF (ImmunoTools, #11343625) |
| Growth media (GM) | SAT10 - bFGF, 1.5 mg/mL Aminocaproic acid (Sigma-Aldrich, #A2504) |
| Differentiation media (DM) | DMEM (Gibco, #11995-065), 2 % Horse serum, 2 mg/mL Aminocarpoic acid, 10 µg/mL insulin (Sigma, #I6634), 1 % Penicillin-Streptomycin |
| Blocking Solution | PBS, 10 % goat serum, 0.3 % Triton X-100 (BioShop, #TRX777) |
| Physiological Salt Solution (PSS) | 140 mM NaCl (Sigma-Aldrich, #S5886), 5mM KCl (Sigma-Aldrich, #P3911), 1mM $MgCl_2$ (Alfa Aesar, #7786-30-3), 10mM HEPES (BioShop, #7365-45-9), 10mM glucose (Sigma-Aldrich, #G8270), 2mM $CaCl_2$ (Sigma-Aldrich, #C1016), corrected to pH 7.3-7.4 |
| Wash Media | 89% DMEM, 10% Fetal Bovine Serum, 1% Penicillin-Streptomycin |

**Supplementary Table 2**

| Antibody | Host Species | Dilution | Manufacturer |
|---|---|---|---|
| DAPI | - | 1:1000 | Roche, #10236276001 |
| Phalloidin 568 | - | 1:400 | Life Technologies, #A12380 |
| Propidium iodide | - | 1:1000 | Sigma-Aldrich, #P4863 |
| DRAQ5 | - | 1:800 | Cell Signaling Technology, #4084L |
| Anti-Sarcomeric α-actinin | Mouse | 1:800 | Sigma-Aldrich, #A7811 |
| Anti-GFP | Chicken | 1:500 | Abcam, #ab13970 |
| Anti-Pax7 | Mouse IgG1 | 1.5:1 | In-house supernatant from hybridoma cell line (DSHB) |
| Anti-Caveolin-1 | Rabbit | 1:300 | Abcam, #ab2910 |
| Anti-c-FOS | Mouse IgG1 | 1:250 | Santa-Cruz, #sc-166940 |
| Anti-Ki67 | Rabbit | 1:300 | Abcam, #ab16667 |
| Anti-N-cadherin | Mouse IgG1 | 1:250 | Santa-Cruz, #sc-8424 |
| Anti-MyoD | Mouse IgG2b | 1:300 | Santa-Cruz, #sc-377460 |
| Anti-CalcR | Rabbit | 1:250 | Abcam, #ab11042 |
| Anti-Integrin α-7 | Rabbit | 1:250 | Abcam, #ab203254 |
| Anti-M-cadherin | Mouse IgG1 | 1:250 | Santa-Cruz, #sc-81471 |
| Anti -Laminin α-2 | Rat | 1:400 | Abcam, #ab11576 |
| Anti-FoxO3a | Mouse IgG1 | 1:250 | Santa-Cruz, #sc-48348 |
| Alexafluor™ 488 Anti-mouse IgG (H+L) | Goat | 1:500 | Invitrogen, #A11001 |
| Alexafluor™ 488 Anti-chicken IgGY (H+L) | Goat | 1:500 | Invitrogen, #A11039 |
| Alexafluor™ 546 Anti-mouse IgG (H+L) | Goat | 1:250 | Invitrogen, #A11003 |

| | | | |
|---|---|---|---|
| Alexafluor™ 546 Anti-rabbit IgG (H+L) | Goat | 1:250 | Invitrogen, #A11010 |
| Alexafluor™ 546 Anti-rat IgG (H+L) | Goat | 1:400 | Invitrogen, #A11081 |
| Alexafluor™ 546 Anti-mouse IgG2b | Goat | 1:300 | Invitrogen, #A21141 |
| Alexafluor™ 555 picolyl azide | - | 1.2:500 | Invitrogen, #C10638B |
| Alexafluor™ 647 Anti-mouse IgG1 | Goat | 1:250 | Invitrogen, #A21240 |
| Alexafluor™ 647 Anti-rabbit IgG (H+L) | Goat | 1:250 | Life Technologies, #A21245 |

**Supplementary Table 3**

| Figure | Independent technical and biological replicates (n, N) | Images per technical replicate (tissue) | n to calculate statistics/error bars | Statistical Test |
|---|---|---|---|---|
| 1D | **SAA coverage** Day 2: n=12 across N=4 Day 5: n=12 across N=4 Day 10: n=15 across N=5 Day 14: n=15 across N=5 Day 16: n=12 across N=4 Day 18: n=12 across N=4 **Fusion Index** Day 2: n=9 across N=3 Day 5: n=12 across N=4 Day 10: n=18 across N=6 Day 14: n=15 across N=5 Day 16: n=6 across N=2 Day 18: n=12 across N=4 | **SAA coverage:** 21 images stitched together **Fusion Index**: 9 | **SAA coverage** Day 2: n=12 Day 5: n=12 Day 10: n=15 Day 14: n=15 Day 16: n=12 Day 18: n=12 **Fusion Index** Day 2: n=9 Day 5: n=12 Day 10: n=18 Day 14: n=15 Day 16: n=6 Day 18: n=12 | One-way ANOVA with Tukey's post-test |
| 1E | Day 2: n=12 across N=4 Day 5: n=12 across N=4 Day 10: n=9 across N=3 Day 14: n=12 across N=4 Day 16: n=12 across N=4 Day 18: n=9 across N=3 | 3 reads | Day 2: n=12 Day 5: n=12 Day 10: n=9 Day 14: n=12 Day 16: n=12 Day 18: n=9 | One-way ANOVA with Tukey's post-test |
| 2D | **200 MuSCs** 1DPE: n=11 across N=4 3DPE: n=12 across N=4 7DPE: n=12 across N=4 **500 MuSCs** 1DPE: n=8 across N=3 3DPE: n=8 across N=3 7DPE: n=9 across N=3 **1500 MuSCs** 1DPE: n=7 across N=3 3DPE: n=9 across N=3 7DPE: n=11 across N=4 **2500 MuSCs** 1DPE: n=9 across N=3 3PE: n=8 across N=3 7DPE: n=9 across N=3 | 25 | **200 MuSCs** 1DPE: n=11 3DPE: n=12 7DPE: n=12 **500 MuSCs** 1DPE: n=8 3DPE: n=8 7DPE: n=9 **1500 MuSCs** 1DPE: n=7 3DPE: n=9 7DPE: n=11 **2500 MuSCs** 1DPE: n=9 3PE: n=8 7DPE: n=9 | One-way ANOVA with Dunnet's test for each individual timepoint comparing against the 500 MuSC condition |

| | | | | |
|---|---|---|---|---|
| 3B | 1DPE: n=9 across N=3 3DPE: n=9 across N=3 7DPE: n=9 across N=3 | 25 | 1DPE: n=9 3DPE: n=9 7DPE: n=9 | One-way ANOVA with Tukey'S post-test comparing the FOS- proportions of each timepoint |
| 3C | 1DPE: n=10 across N=3 3DPE: n=11 across N=4 7DPE: n=11 across N=4 | 25 | 1DPE: n=10 3DPE: n=11 7DPE: n=11 | One-way ANOVA with Tukey's post-test comparing the Ki67- proportions of each timepoint |
| 3E | n=15 across N=5 | 25 | n=15 | - |
| 3G | **PSS** n=16 across N=5 **2.4% BaCl** n=18 across N=6 | 25 | **PSS** n=16 **2.4% BaCl** n=18 | Unpaired t-test of the Ki67- proportions of both conditions |
| 4B | 1DPE: n=6 across N=2 3DPE: n=7 across N=2 7DPE: n=6 across N=2 | 25 | 1DPE: n=6 3DPE: n=7 7DPE: n=6 | - |
| 4C | 1DPE: n=9 across N=3 3DPE: n=9 across N=3 7DPE: n=6 across N=2 | 25 | 1DPE: n=9 3DPE: n=9 7DPE: n=6 | - |
| 4D | 1DPE: n=6 across N=2 3DPE: n=6 across N=2 7DPE: n=6 across N=2 | 25 | 1DPE: n=6 3DPE: n=6 7DPE: n=6 | - |
| 4E | 1DPE: n=6 across N=2 3DPE: n=6 across N=2 7DPE: n=6 across N=2 | 25 | 1DPE: n=6 3DPE: n=6 7DPE: n=6 | - |
| 4F | 1DPE: n=10 across N=3 3DPE: n=8 across N=3 7DPE: n=8 across N=3 | 25 | 1DPE: n=10 3DPE: n=8 7DPE: n=8 | - |
| 5C | 1DPE: n=916 across N=4 3DPE: n=980 across N=4 7DPE: n=737 across N=3 | 25 | - | - |

| | | | | |
|---|---|---|---|---|
| 7A | **Young** 1DPE: n=9 across N=3 3DPE: n=9 across N=3 7DPE: n=9 across N=3 **Young+Wortmannin** 1DPE: n=6 across N=2 3DPE: n=6 across N=2 7DPE: n=6 across N=2 **Aged** 1DPE: n=9 across N=3 3DPE: n=8 across N=3 7DPE: n=9 across N=3 **Aged+Wortmannin** 1DPE: n=9 across N=3 3DPE: n=9 across N=3 7DPE: n=7 across N=3 | 25 | **Young** 1DPE: n=9 3DPE: n=9 7DPE: n=9 **Young+Wortmannin** 1DPE: n=6 3DPE: n=6 7DPE: n=6 **Aged** 1DPE: n=9 3DPE: n=8 7DPE: n=9 **Aged+Wortmannin** 1DPE: n=9 3DPE: n=9 7DPE: n=7 | One-way ANOVA with Dunnet's test for each individual timepoint comparing against the Young condition |
| 7C | **Young** 1DPE: n=9 across N=3 3DPE: n=9 across N=3 7DPE: n=9 across N=3 **Young+Wortmannin** 1DPE: n=6 across N=2 3DPE: n=6 across N=2 7DPE: n=6 across N=2 **Aged** 1DPE: n=9 across N=3 3DPE: n=8 across N=3 7DPE: n=9 across N=3 **Aged+Wortmannin** 1DPE: n=9 across N=3 3DPE: n=9 across N=3 7DPE: n=7 across N=3 | 25 | **Young** 1DPE: n=9 3DPE: n=9 7DPE: n=9 **Young+Wortmannin** 1DPE: n=6 3DPE: n=6 7DPE: n=6 **Aged** 1DPE: n=9 3DPE: n=8 7DPE: n=9 **Aged+Wortmannin** 1DPE: n=9 3DPE: n=9 7DPE: n=7 | One-way ANOVA with Tukey’s post-test comparing the conditions against each other at the 3 DPE timepoint |
| 7D | **Young** 1DPE: n=6 across N=2 3DPE: n=6 across N=2 7DPE: n=5 across N=2 **Young+Wortmannin** 1DPE: n=5 across N=2 3DPE: n=5 across N=2 7DPE: n=5 across N=2 **Aged** 1DPE: n=5 across N=2 3DPE: n=8 across N=3 7DPE: n=10 across N=3 **Aged+Wortmannin** 1DPE: n=5 across N=2 3DPE: n=6 across N=2 7DPE: n=6 across N=2 | 25 | **Young** 1DPE: n=6 3DPE: n=6 7DPE: n=5 **Young+Wortmannin** 1DPE: n=5 3DPE: n=5 7DPE: n=5 **Aged** 1DPE: n=5 3DPE: n=8 7DPE: n=10 **Aged+Wortmannin** 1DPE: n=5 3DPE: n=6 7DPE: n=6 | One-way ANOVA with Tukey’s post-test comparing the conditions against each other at the 3 DPE timepoint |
| 7E | **Young** n=9 across N=3 **Young+Wortmannin** n=6 across N=2 **Aged** n=9 across N=3 **Aged+Wortmannin** n=9 across N=3 | 25 | - | - |
| 7F | **Young** n=9 across N=3 **Young+Wortmannin** n=6 across N=2 **Aged** n=9 across N=3 **Aged+Wortmannin** n=9 across N=3 | 25 | **Young** n=9 **Young+Wortmannin** n=6 **Aged** n=9 **Aged+Wortmannin** n=9 | One-way ANOVA with Tukey’s post-test |

| | | | | |
|---|---|---|---|---|
| 7G | **Young** n=9 across N=3 **Young+Wortmannin** n=6 across N=2 **Aged** n=9 across N=3 **Aged+Wortmannin** n=9 across N=3 | 25 | **Young** n=9 **Young+Wortmannin** n=6 **Aged** n=9 **Aged+Wortmannin** n=9 | One-way ANOVA with Tukey's post-test |
| S1B | **10,000** n=12 across N=4 **25,000** n=12 across N=4 **50,000** n=12 across N=4 | 21 images stitched together | **10,000** n=12 **25,000** n=12 **50,000** n=12 | One-way ANOVA with Tukey's post-test |
| S2C | n=11 across N=4 | 25 | n=11 | - |
| S3B | 1DPE: n=8 across N=3 3DPE: n=7 across N=3 7DPE: n=15 across N=5 | 25 | 1DPE: n=8 3DPE: n=7 7DPE: n=15 | One-way ANOVA with Tukey's post-test |
| S4A | **BI** n=8 across N=3 **PSS** n=7 across N=3 **2.4% BaCl** n=9 across N=3 | 21 images stitched together | **BI** n=8 **PSS** n=7 **2.4% BaCl** n=9 | One-way ANOVA with Tukey's post-test |
| S4B | **BI** n=7 across N=3 **PSS** n=9 across N=3 **2.4% BaCl** n=9 across N=3 | 25 | **BI** n=7 **PSS** n=9 **2.4% BaCl** n=9 | One-way ANOVA with Tukey's post-test |
| S5C | **200 MuSCs** n=11 across N=4 **500 MuSCs** n=15 across N=5 **1500 MuSCs** n=16 across N=5 **2500 MuSCs** n=13 across N=4 | 25 | **200 MuSCs** n=11 **500 MuSCs** n=15 **1500 MuSCs** n=16 **2500 MuSCs** n=13 | One-way ANOVA with Tukey's post-test |
| S5D | **200 MuSCs** n=12 across N=4 **500 MuSCs** n=9 across N=3 **1500 MuSCs** n=12 across N=4 **2500 MuSCs** n=9 across N=4 | 25 | **200 MuSCs** n=12 **500 MuSCs** n=9 **1500 MuSCs** n=12 **2500 MuSCs** n=9 | One-way ANOVA with Tukey's post-test |
| S6B | **Day 5** n=15 across N=5 **Day 0** n=11 across N=4 | 25 | **Day 5** n=15 5 **Day 0** n=11 | Unpaired t-test |
| S8C | 1DPE: n=35 across N=3 3DPE: n=45 across N=3 7DPE: n=45 across N=3 | Every 5 images is from 1 tissue | n=125 | Simpe linear regression |
| S9A | 1DPE: n=916 across N=4 3DPE: n=980 across N=4 7DPE: n=737 across N=3 | 25 | 1DPE: n=916 3DPE: n=980 7DPE: n=737 | - |

| | | | | |
|---|---|---|---|---|
| S9C | **Pax7+/MyoD-** 3DPE: n=12 across N=4 7DPE: n=11 across N=3 **Pax7+/MyoD+** 3DPE: n=11 across N=4 7DPE: n=10 across N=3 | 25 | **Pax7+/MyoD-** 3DPE: n=12 7DPE: n=11 **Pax7+/MyoD+** 3DPE: n=11 7DPE: n=10 | Unpaired t-tests |
| S12A | **Young** Day 1: n=6 across N=2 Day 3: n=6 across N=2 **Young+Wortmannin** Day 1: n=6 across N=2 Day 3: n=6 across N=2 **Aged** Day 1: n=6 across N=2 Day 3: n=6 across N=2 **Aged+Wortmannin** Day 1: n=6 across N=2 Day 3: n=6 across N=2 | 104 | **Young** Day 1: n=6 Day 3: n=6 **Young+Wortmannin** Day 1: n=6 Day 3: n=6 **Aged** Day 1: n=6 Day 3: n=6 **Aged+Wortmannin** Day 1: n=6 Day 3: n=6 | One-way ANOVA with Tukey’s post-test comparing each experimental group at the 3 DPE timepoint |
| S12B | **Young** n=2716 across N=2 **Young+Wortmannin** n=565 across N=2 **Aged** n=4437 across N=2 **Aged+Wortmannin** n=1897 across N=2 | 104 | **Young** n=2716 **Young+Wortmannin** n=565 **Aged** n=4437 **Aged+Wortmannin** n=1897 | Outliers removed with the ROUT method (with Q=1%) and one-way ANOVA performed with Šidák’s post-test comparing pre-selected conditions |
| S13B | **Young** n=9 across N=3 **Young+Wortmannin** n=6 across N=2 **Aged** n=8 across N=3 **Aged+Wortmannin** n=7 across N=3 | 25 | **Young** n=9 **Young+Wortmannin** n=6 **Aged** n=8 **Aged+Wortmannin** n=7 | One-way ANOVA with Tukey's post-test |
